# Type V Collagen Controls Decidual Extracellular Matrix Organization and Angiogenesis During Embryo Implantation

**DOI:** 10.64898/2026.07.22.740039

**Authors:** Mona Gebril, Emily Kinneston, Rimi Das, Joseph Boyd, John P Lydon, Shanmugasundaram Nallasamy

**Author notes:** **Corresponding author:** Shanmugasundaram Nallasamy, DVM, PhD Assistant Professor, Division of Reproductive Sciences, Dept. of Obstetrics, Gynecology and Reproductive Sciences University of Vermont College of Medicine, 95 Carrigan Drive, Burlington, VT 05405.

## Abstract

Extracellular matrix (ECM) remodeling and angiogenesis are essential processes underlying endometrial decidualization during embryo implantation. Fibrillar collagens, the major structural components of the ECM, form the architectural framework of the decidua and undergo dynamic reorganization during this process. However, the functional role of type V collagen—a key regulator of collagen fibrillogenesis— remains undefined. Here, we demonstrate that *Col5a1* is highly expressed in decidual stromal and endothelial cells of the mouse uterus. Conditional deletion of *Col5a1* leads to progressive uterine hemorrhage beginning at gestation day 8, culminating in complete embryo resorption and pregnancy loss by day 12. *Col5a1*-deficient decidua exhibits severe structural distortion, marked disorganization of fibrillar collagen, shallow and misdirected embryo invasion, and profound disruption of decidual angiogenesis and vascular network formation. Transcriptomic profiling further reveals distinct gene expression signatures and signaling pathways regulated by COL5A1 in the decidua. Collectively, these findings identify type V collagen as a critical ECM regulator required for maintaining decidual integrity, supporting angiogenesis, and ensuring successful pregnancy.

## INTRODUCTION

The majority of early pregnancy losses occur during the peri- and early post-implantation periods of the first trimester of pregnancy^1–3^. Defects in implantation-specific events, such as decidualization and angiogenesis, are also associated with later pregnancy complications, including placental dysfunction, intrauterine growth restriction, preeclampsia, and recurrent pregnancy loss^3–7^. Thus, elucidating the molecular and cellular mechanisms that drive decidualization and angiogenesis will help identify targeted approaches to improve pregnancy outcomes. Attachment of the embryo to the receptive endometrium triggers localized stromal cell proliferation and differentiation, a process known as endometrial decidualization^8,9^. Decidualization is a critical event during early pregnancy, resulting in the formation of a specialized tissue structure, the “decidua,” which supports the embryo until the establishment of a mature placenta. In parallel with these stromal cell functions, key underlying processes that support decidual function include uterine extracellular matrix (ECM) reorganization and angiogenesis^10–12^.

ECM reorganization, defined as alterations in structural composition and organization, is a concomitant process of decidualization^13,14^. Fibrillar collagens are the most abundant group of ECM proteins and provide structural integrity while regulating multiple cellular functions in various tissues^15,16^. During endometrial decidualization, fibrillar collagens undergo dynamic remodeling, resulting in significant changes in their expression, spatial distribution, and structural organization within the mouse uterine decidua^13^. These fibrillar collagens reorient and align parallel to the direction of embryo invasion, thereby facilitating this process^13^. Furthermore, genes encoding factors involved in the synthesis, processing, and assembly of fibrillar collagen exhibit differential expression during decidualization^13^. Uterine angiogenesis establishes the decidual vascular network, which not only nourishes the invading embryo but also supports the development of the decidua and placenta^11^.

Type V collagen, a minor fibrillar collagen but a key regulator of collagen fibrillogenesis, forms heterotypic fibrils with the predominant type I collagen to generate mature fibrillar collagen^17–20^. It is indispensable for the nucleation and assembly of collagen fibrils. It consists of two α1(V) chains and one α2(V) chain and is encoded by the *Col5a1*, *Col5a2*, and *Col5a3* genes. Notably, *Col5a1* (collagen type V alpha 1 chain) knockout mice are embryonic lethal due to failure of fibrillar collagen synthesis and assembly^20^. Furthermore, conditional deletion of *Col5a1* leads to dysregulation of collagen fibrillogenesis in the cornea, tendon, and ligament^18,19^. Mutations in collagen V are highly prevalent in patients with classic Ehlers–Danlos syndrome (EDS) of which approximately 75% occur in *COL5A1*^21,22^. Thus, loss or mutation of type V collagen results in defective fibrillar collagen synthesis and assembly, leading to compromised tissue function. Collectively, these findings establish type V collagen as a critical regulator of collagen fibrillogenesis, matrix assembly, and tissue integrity. In addition to connective tissues, type V collagen is highly expressed in endothelial cells and the vascular ECM of many tissues^22–24^. *Col5a1* zebrafish mutants exhibit spontaneous trunk hemorrhage due to compromised vascular integrity and fail to survive^25^. Heterozygous *Col5a1* mice display characteristics of EDS and exhibit blood vessels with reduced stiffness and strength^20^. In humans, *COL5A1* mutations are associated with arterial dissection^26,27^. Collectively, these studies demonstrate a critical role for type V collagen in maintaining vascular integrity and function, which is a prerequisite for angiogenesis.

Type V collagen is expressed in nearly all tissues of the female reproductive tract, including the uterus^10,14^. However, the functional role of type V collagen in endometrial decidualization, embryo invasion, and angiogenesis remains unknown. In this study, we provide compelling evidence, using a mouse model, supporting the involvement of type V collagen in decidualization and angiogenesis. Conditional deletion of *Col5a1* results in complete pregnancy loss in mice due to severe decidual defects in ECM reorganization and disruption of the decidual vascular network. These findings identify type V collagen as a critical ECM regulator essential for maintaining decidual integrity, supporting angiogenesis, and ensuring successful pregnancy.

## RESULTS

### Uterine type V collagen is essential for successful pregnancy in mice

We conducted a comprehensive analysis of *Col5a1* expression in the uterus during embryo implantation (Supplementary Fig. S1). RNAscope analysis revealed that *Col5a1* transcripts are widely distributed throughout the decidua from gestation day (GD) 6 to 8. Notably, expression was particularly abundant within the secondary decidual zone and in stromal cells adjacent to the myometrium (Supplementary Fig. S1A). Consistent with the RNA localization pattern, COL5A1 protein was highly expressed throughout the decidua during GD6 to 9, with a relatively uniform distribution in both mesometrial and antimesometrial regions (Supplementary Fig. S1B). To determine the functional role of type V collagen during embryo implantation, we generated *Col5a1* conditional knockout (*Col5a1^d/d^*) mice. Efficient deletion of *Col5a1* in the uterus was confirmed by quantitative real-time PCR and RNAscope analysis. As shown in Supplementary Fig. S2, *Col5a1* expression was markedly reduced in the decidua of *Col5a1^d/d^* mice compared with *Col5a1^f/f^* mice, thereby confirming the successful deletion of *Col5a1* from the uterus. To assess fertility, we conducted a six-month breeding study. As summarized in supplementary table 1, all *Col5a1^f/f^* females delivered pups, whereas none of the *Col5a1^d/d^* females produced offspring, demonstrating that *Col5a1* is essential for female fertility.

To investigate the cause of fertility defects in *Col5a1^d/d^* female mice, we first assessed ovarian function by inducing superovulation. Prepubertal *Col5a1^f/f^* and *Col5a1^d/d^* mice were treated with a standard gonadotropin regimen. Following hormonal stimulation, the number of ova recovered from *Col5a1^d/d^* females was comparable to that from *Col5a1^f/f^* controls (Supplementary Fig. S3A, B), indicating that ovulation is not impaired by loss of *Col5a1*. Consistent with normal ovarian function, serum progesterone and estrogen levels measured on GD6 were similar between *Col5a1^f/f^* and *Col5a1^d/d^* females (Supplementary Fig. S3C). Together, these findings suggest that pregnancy loss in *Col5a1^d/d^* females is not attributable to dysfunction of the hypothalamic–pituitary– ovarian axis. To further investigate the cause of fertility defects, gravid uteri from timed-pregnant *Col5a1^d/d^* females were examined. Gross morphological analysis revealed no discernible differences in implantation sites between *Col5a1^f/f^* and *Col5a1^d/d^* uteri on GD6, indicating that events preceding GD6 are similar between these two genotypes (Fig. 1A). However, by GD8, *Col5a1^d/d^* uteri displayed pronounced uterine hemorrhage, which progressively worsened over time and culminated in embryo resorption and complete pregnancy loss by GD12 (Fig. 1A). The number of healthy implantation sites was progressively and significantly reduced, whereas the number of resorption sites progressively increased in *Col5a1^d/d^* uteri compared to *Col5a1^f/f^* uteri (Fig. 1B). Collectively, these findings demonstrate that uterine deletion of *Col5a1* results in complete pregnancy loss in mice, demonstrating that type V collagen is necessary for successful pregnancy.

**Fig. 1.**
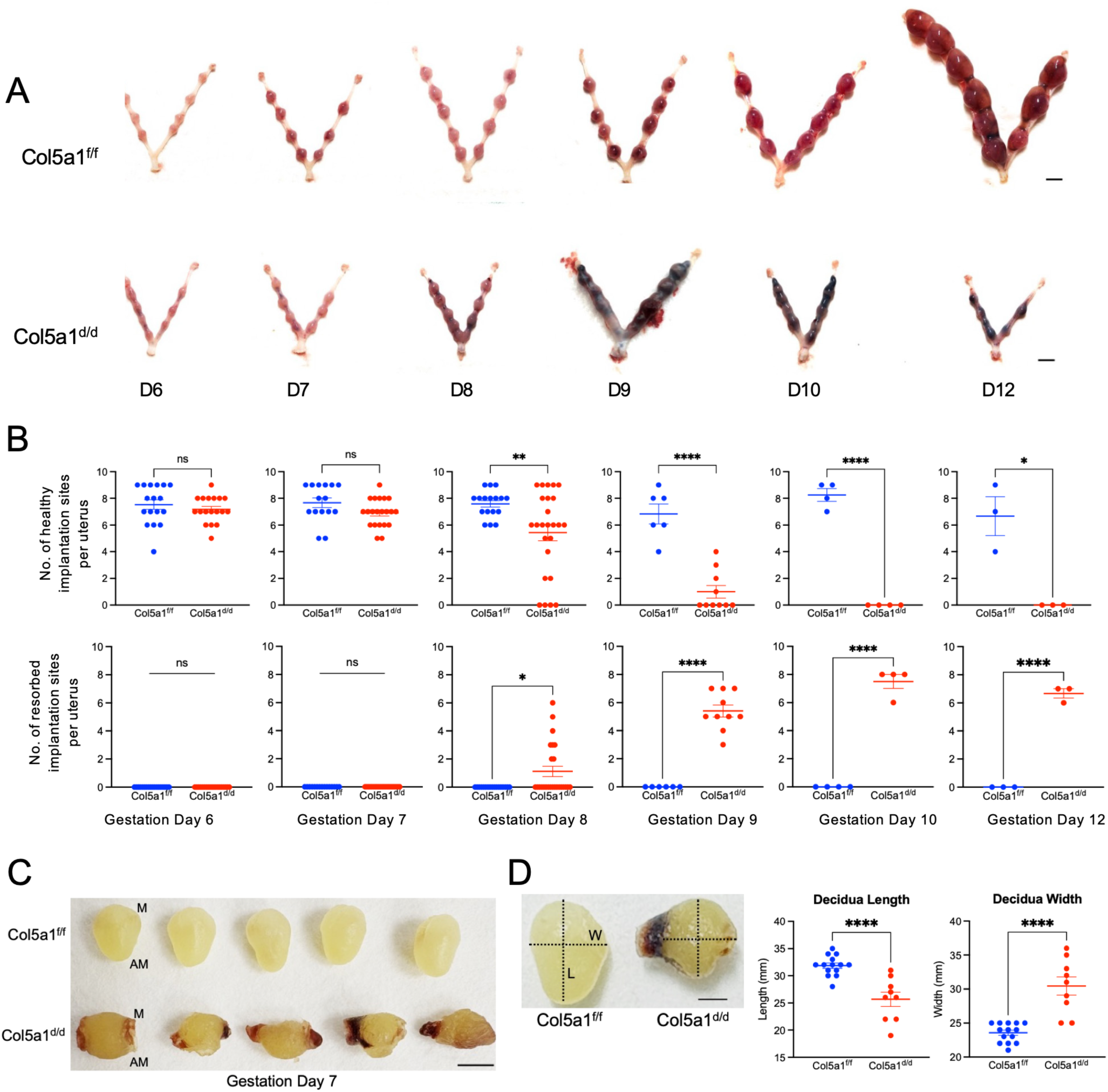
Conditional deletion of *Col5a1* causes complete pregnancy loss in mice. (A) Evaluation of gross morphology of pregnant uteri from *Col5a1^f/f^* and *Col5a1^d/d^* mice from gestation day (GD) 6 to GD12. (B) Quantification of healthy and resorbed implantation sites at specific gestational time points. The number of uteri examined, and the numbers of healthy and resorbed implantation sites measured, are provided in Supplementary Table 2. (C) Evaluation of decidua dissected from uteri of *Col5a1^f/f^* and *Col5a1^d/d^* mice on GD7 (n = 3 mice per genotype). (D) Length and width measurements of decidua collected from *Col5a1^f/f^* and *Col5a1^d/d^* mice on GD7 (n = 3 mice per genotype). M - mesometrial region, AM - antimesometrial region, W – Width, L – Length. ns, not significant; *P < 0.05; **P < 0.01; ****P < 0.0001.

### Type V collagen is necessary for the preservation of decidual tissue integrity

A comparative analysis of dissected decidua from *Col5a1^f/f^* and *Col5a1^d/d^*mice at GD7 revealed notable disparities in their morphology. The *Col5a1^f/f^* decidua were acorn-shaped, with a broader mesometrial end and a narrowed antimesometrial end. In contrast, the *Col5a1^d/d^* decidua displayed signs of bleeding, were highly fragile, bean-shaped, and longitudinally embedded within the uterus (Fig. 1C). Consistent with these morphological distortions, both length and width were significantly altered, suggesting loss of decidual tissue integrity (Fig. 1D). Histological examination further demonstrated striking differences in the morphology of uterine sections at the implantation site. In *Col5a1^f/f^* controls, uterine sections appeared elongated and exhibited temporal growth and size characteristics appropriate for mouse implantation site development. In contrast, *Col5a1^d/d^* uterine sections appeared rounded and smaller compared to the larger, elongated sections observed in *Col5a1^f/f^* mice (Fig. 2A).

**Fig. 2.**
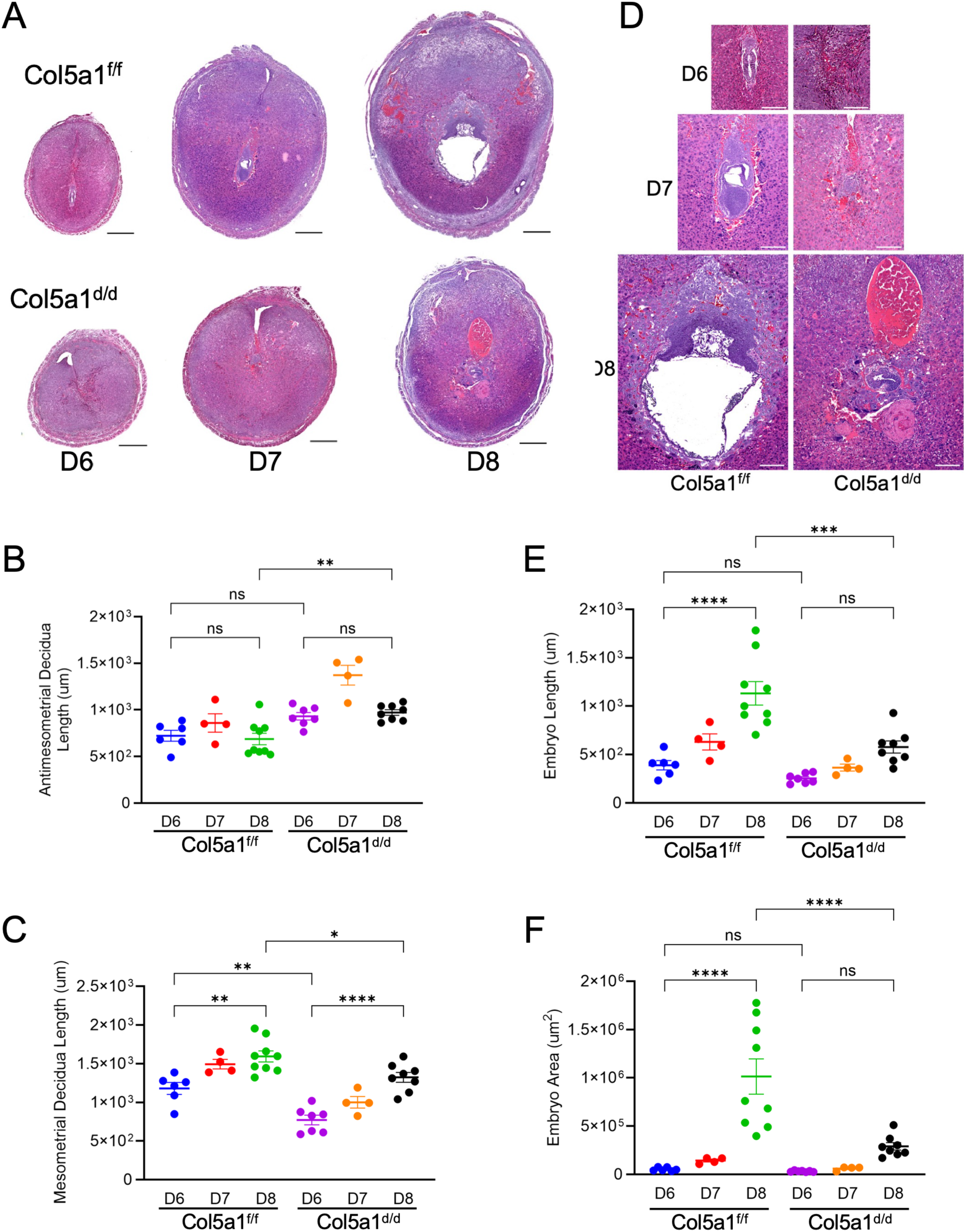
Impaired embryo invasion in uteri lacking *Col5a1*. (A) Histopathological evaluation of uterine sections at implantation sites prepared from *Col5a1^f/f^* and *Col5a1^d/d^* mice at GD6– GD8 stained with H&E. Scale bar, 500 µm. (B) Quantitative analysis of antimesometrial decidual length at GD6–GD8 using uterine sections from *Col5a1^f/f^* and *Col5a1^d/d^* mice. (C) Quantitative analysis of mesometrial decidual length at GD6–GD8 using uterine sections from *Col5a1^f/f^* and *Col5a1^d/d^* mice. (D) Uterine sections at implantation sites displaying embryos at GD6–GD8. Scale bar, 200 µm. (E) Quantitative analysis of embryo length at GD6–GD8 using uterine sections from *Col5a1^f/f^* and *Col5a1^d/d^* mice. (F) Quantitative analysis of embryo area at GD6–GD8 using uterine sections from *Col5a1^f/f^* and *Col5a1^d/d^* mice (n = 4–9 mice per gestational time point per genotype). The measurement guide is shown in Supplementary Fig. 4. ns, not significant; *P < 0.05; **P < 0.01; ***P < 0.001; ****P < 0.0001.

We measured the length of antimesometrial and mesometrial decidua as described in Supplementary Fig. 4. The length of the antimesometrial decidua (defined as the distance from the distal end of the embryo to the end of the antimesometrial decidua at the myometrium) at GD6 did not differ significantly between *Col5a1^f/f^* and *Col5a1^d/d^* mice. However, the antimesometrial distance was significantly higher at GD8 in *Col5a1^d/d^* mice compared to *Col5a1^f/f^* mice (Fig. 2B). When examining the mesometrial decidua (defined as the distance from the inner myometrium to the embryo on the mesometrial side), this distance increased progressively from GD6 through GD8 in *Col5a1^f/f^* uteri, reflecting expansion of the decidua at the mesometrial side, the site destined for future placentation. Although mesometrial decidual length increased progressively in *Col5a1^d/d^* uteri, the length was significantly reduced compared to *Col5a1^f/f^* uteri (Fig. 2C). These findings are consistent with our previous observations, including distorted decidual size and shape, and collectively demonstrate that type V collagen is necessary to maintain decidual tissue integrity for successful embryo implantation.

### Type V collagen provides directional cues for embryo invasion within the decidua

Our histological evaluation demonstrated that embryos invaded the antimesometrial decidua and exhibited stage-specific development in *Col5a1^f/f^* mice (Fig. 2A–C). However, embryos failed to invade deeper into the antimesometrial decidua in *Col5a1^d/d^* mice (Fig. 2A–C). Notably, embryo orientation was markedly altered, appearing completely tilted and aligned horizontally relative to the mesometrial–antimesometrial axis at GD8 in *Col5a1^d/d^* mice (Fig. 2D). Both embryo length and area were significantly reduced in *Col5a1^d/d^* decidua (Fig. 2E, F). To further examine embryo morphology within the decidua, trophoblast cells were visualized by pan-keratin staining. In *Col5a1^f/f^* uteri, embryos exhibited stage-specific growth and development and maintained proper mesometrial–antimesometrial orientation during invasion of the decidua. In contrast, embryo orientation within the *Col5a1^d/d^* decidua was abnormal. Embryo invasion in *Col5a1^d/d^* uteri was severely disrupted, with embryos exhibiting a tilted mesometrial–antimesometrial axis, perpendicular to the normal axis of invasion (Fig. 3A).

**Fig. 3.**
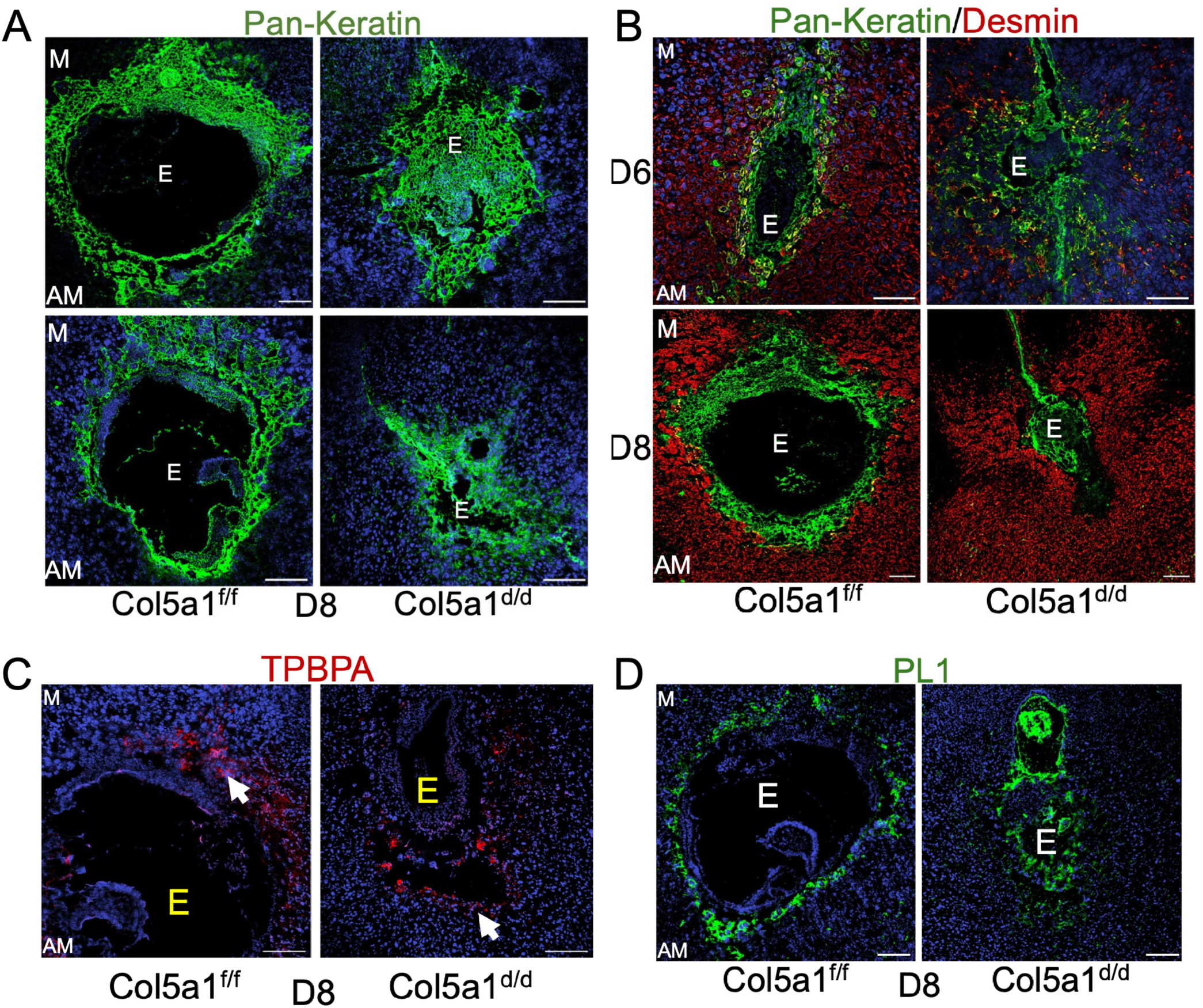
Embryo orientation and invasion direction are compromised in *Col5a1*-deficient decidua. Visualization of embryo orientation and invasion direction within *Col5a1^f/f^* and *Col5a1^d/d^*decidua. (A) Confocal imaging of pan-keratin in frozen uterine sections from *Col5a1^f/f^* and *Col5a1^d/d^* mice on GD8. (B) Confocal imaging of pan-keratin (green) and desmin (red) in frozen uterine sections from *Col5a1^f/f^* and *Col5a1^d/d^* mice on GD6 and GD8. (C) Confocal imaging of trophoblast-specific protein α (TPBPA; red) in frozen uterine sections from *Col5a1^f/f^* and *Col5a1^d/d^* mice on GD8. (D) Confocal imaging of placental lactogen-1 (PL-1; green) in frozen uterine sections from *Col5a1^f/f^* and *Col5a1^d/d^* mice on GD8. Image acquisition and analysis settings were adjusted for each image to optimize morphological visualization. AM, images captured from the antimesometrial region; M, images captured from the mesometrial region. Representative images from three independent experiments. DAPI (blue). Scale bar, 200 µm.

**Fig. 4.**
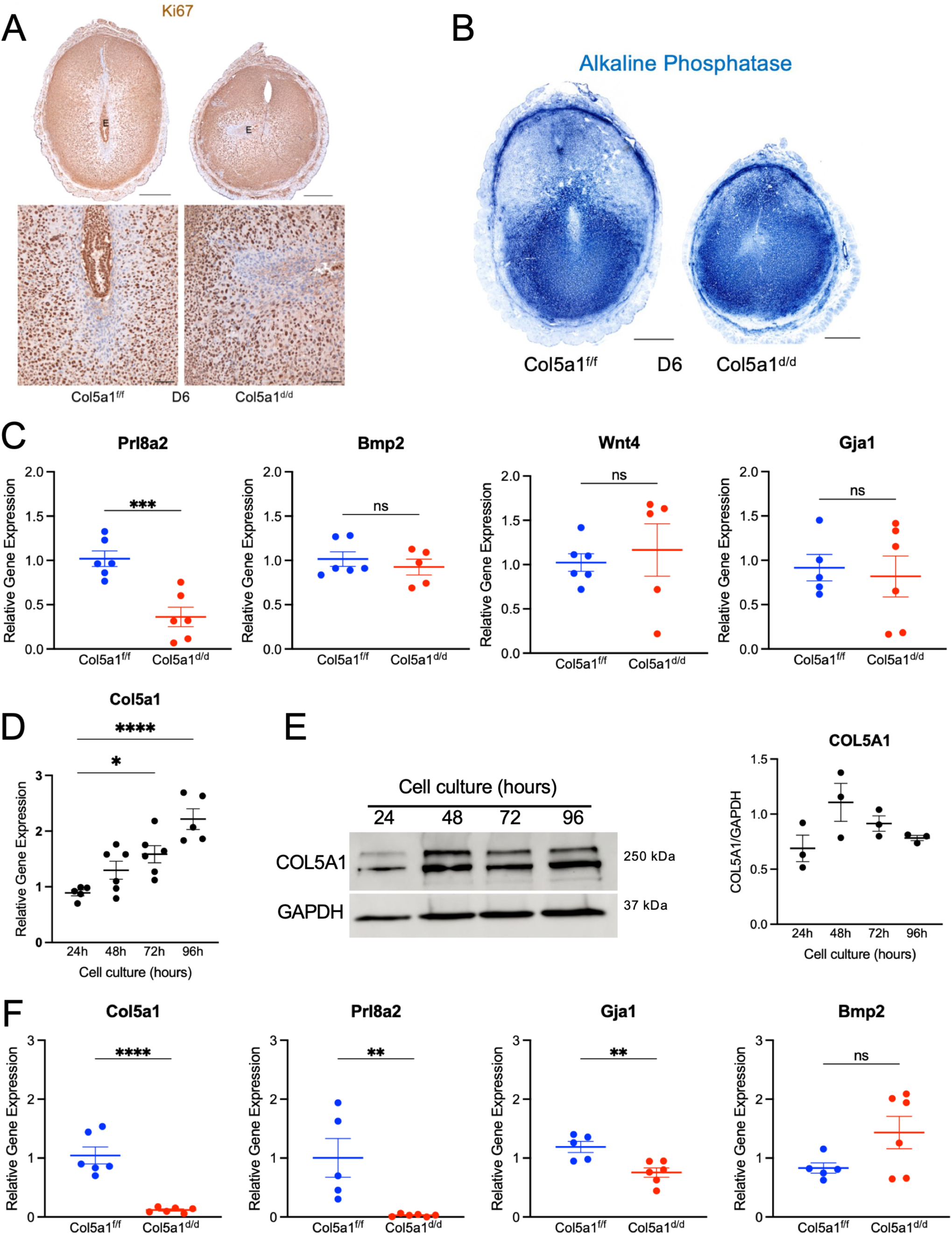
Selective impairment of decidualization in *Col5a1^d/d^* uteri. (A) Immunohistochemical localization of Ki67 in uterine sections from *Col5a1^f/f^* and *Col5a1^d/d^* mice on GD6. Scale bar, 500 µm (upper panel) and 100 µm (lower panel) (n = 3 mice per genotype). (B) Alkaline phosphatase activity in uterine sections from *Col5a1^f/f^* and *Col5a1^d/d^* mice on GD6. Scale bar, 500 µm (n = 6 mice per genotype). (C) Gene expression analysis of decidualization markers (*Prl8a2*, *Bmp2*, *Wnt4*, and *Gja1*) using total RNA isolated from GD6 decidual tissues of *Col5a1^f/f^* and *Col5a1^d/d^* mice. Gene expression was normalized to *Rplp0* (n = 5–6 mice per group). (D) Gene expression of *Col5a1* in mouse endometrial stromal cells undergoing in vitro decidualization. Cells isolated from GD3 uteri were cultured, and lysates were collected at 24, 48, 72, and 96 h for RNA analysis. Gene expression was normalized to *Rplp0* and compared with 24 h samples (n = 5–6 per group). (E) Western blot analysis of COL5A1 protein levels in cell lysates collected at 24, 48, 72, and 96 h. GAPDH was used as a loading control. Representative images from three independent experiments are shown. Quantification of COL5A1 protein levels is presented in the graph. (F) Gene expression of *Col5a1*, *Prl8a2*, *Gja1*, and *Bmp2* in mouse endometrial stromal cells undergoing in vitro decidualization. Cells isolated from GD3 *Col5a1^f/f^* and *Col5a1^d/d^* uteri were cultured, and lysates were collected at 96 h for RNA analysis. Gene expression was normalized to *Rplp0* (n = 5–6 per group). ns, not significant; *P < 0.05; **P < 0.01; ***P < 0.001; ****P < 0.0001.

To further evaluate trophoblast invasion within the decidua, decidual cells were stained with desmin and trophoblast cells with pan-keratin. In *Col5a1^f/f^* uteri, trophoblast cells were tightly associated with decidual tissues at GD6 and GD8. In contrast, embryos in *Col5a1^d/d^* uteri were loosely associated with decidual tissues and exhibited abnormal orientation. The shape and size of embryos within implantation sites were also abnormal (Fig. 3B). To further assess trophoblast growth and expansion, we examined the spatial distribution of spongiotrophoblast cells using the trophoblast-specific marker trophoblast-specific protein alpha (TPBPA). In *Col5a1^f/f^* embryos, TPBPA expression was localized to the ectoplacental cone, whereas in *Col5a1^d/d^* embryos, TPBPA expression was aberrantly detected within the antimesometrial decidua (Fig. 3C). Localization of placental lactogen 1 (PL-1), a trophoblast giant cell-specific marker, further revealed disrupted and structurally compromised trophoblast cells within the *Col5a1^d/d^* decidua (Fig. 3D). Collectively, these results demonstrate that type V collagen–mediated decidual integrity is essential for proper embryonic growth and invasion, as well as for maintaining appropriate mesometrial– antimesometrial orientation within the decidua.

### Type V collagen regulates specific pathways involved in decidualization

Our findings revealed that, in *Col5a1^d/d^* uteri on GD6, embryos were able to attach and invade into the stromal compartment, indicating that critical events preceding this time point, such as uterine receptivity, were similar between *Col5a1^f/f^* and *Col5a1^d/d^* mice. Consistent with these observations, uterine expression of progesterone receptor (PGR) and estrogen receptor alpha (ESR1) in *Col5a1^d/d^* mice was comparable to that observed in *Col5a1^f/f^* controls (Supplementary Fig. 5). Next, we examined stromal cell proliferation by localizing Ki67, a marker of cell proliferation. The expression levels of Ki67 in GD6 uterine sections were comparable between *Col5a1^f/f^* and *Col5a1^d/d^* mice (Fig. 4A). Next, we examined stromal cell decidualization. Although the levels of alkaline phosphatase (ALP), a marker of decidualization, were comparable between *Col5a1^f/f^* and *Col5a1^d/d^* uterine sections, the pattern of ALP staining differed between the two groups. In *Col5a1^f/f^* uteri, ALP staining was more intense in the antimesometrial decidua compared to the mesometrial decidua. In contrast, in *Col5a1^d/d^* uterine sections, ALP staining lost its region-specific pattern and was diffusely distributed throughout the decidua (Fig. 4B).

**Fig. 5.**
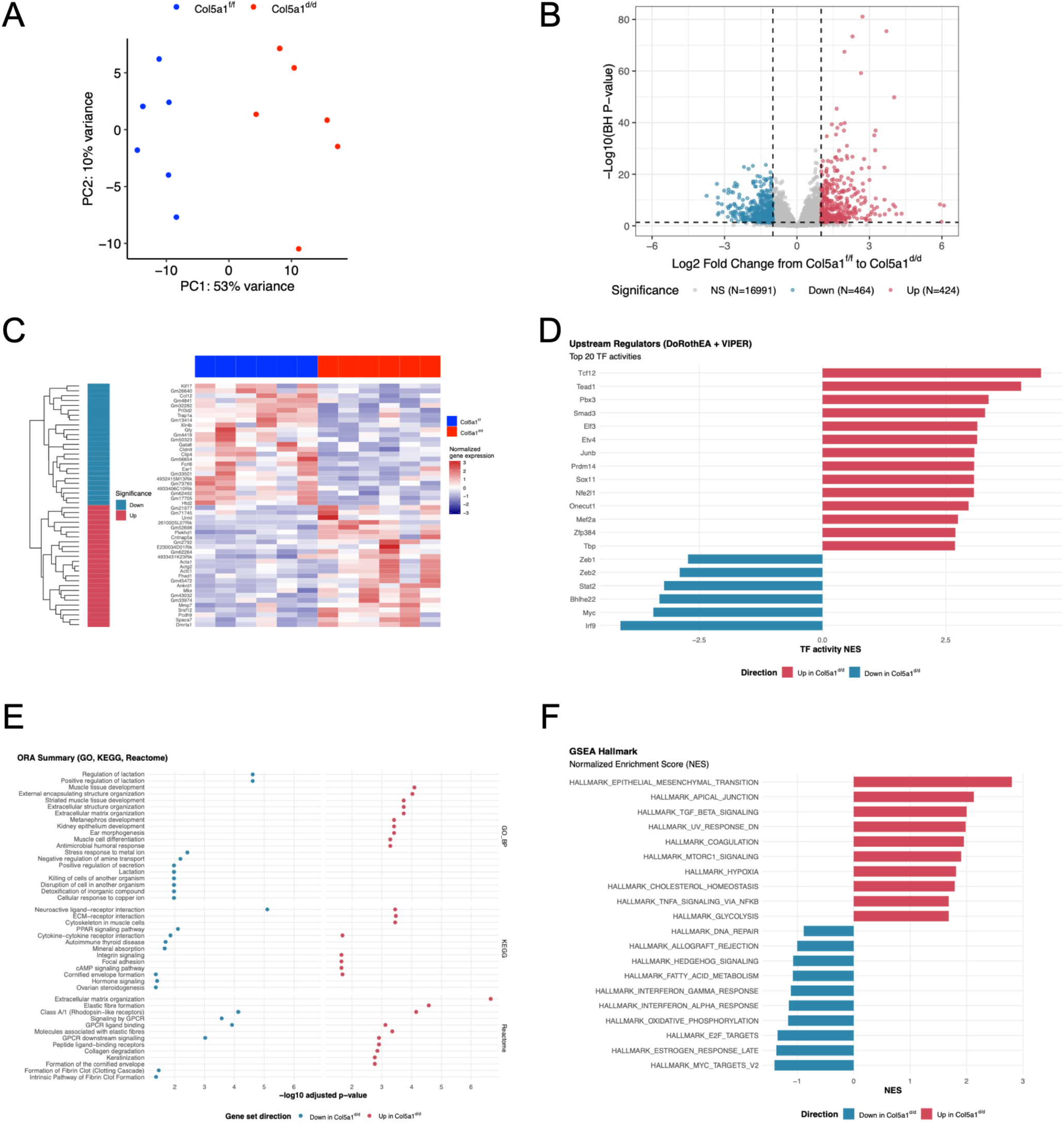
Altered transcriptomic profiling in *Col5a1^d/d^* decidua as determined by RNA sequencing analysis. (A) Principal component analysis of RNA-seq data from *Col5a1^f/f^* and *Col5a1^d/d^* decidual samples collected on GD6. Read counts were normalized using the rlog function from DESeq2. (B) Volcano plot summarizing differential gene expression by fold change and statistical significance. Genes were considered significant when |log₂ fold change| > 1 and adjusted *P* < 0.05. Genes with the greatest magnitude of change and statistical significance are located furthest from the center and highest on the plot. (C) Heatmap showing expression patterns across selected genes and samples, highlighting coordinated transcriptional differences between groups. (D) Plot ranking transcription factors by inferred normalized enrichment scores. The most activated and most repressed regulators are at the extremes, whereas transcription factors near zero show minimal coherent signal across their targets. (E) Over-representation analysis (ORA) dot plot summarizing enriched functional categories among differentially expressed genes. (F) Plot summarizing Hallmark pathway enrichment scores. Positive and negative normalized enrichment scores indicate opposite directions of regulation.

Prolactin family 8, subfamily a, member 2 (*Prl8a2*) is a well-established marker of stromal cell decidualization in mice and is significantly induced during the differentiation of endometrial stromal cells into decidual cells^28^. Notably, *Prl8a2* expression was significantly reduced in *Col5a1^d/d^* decidua compared to *Col5a1^f/f^* decidua. However, the expression of other well-known regulators of stromal decidualization, such as *Bmp2*, *Wnt4*, and *Gja1*, was not altered in *Col5a1^d/d^* mice (Fig. 4C). To specifically examine stromal cell decidualization, we utilized an in vitro stromal cell decidualization model. *Col5a1* mRNA was significantly induced during in vitro stromal cell decidualization (Fig. 4D). The protein levels of COL5A1 were abundant at all time points analyzed, but the levels remained relatively constant across all time points (Fig. 4E). The expression of *Col5a1* was significantly reduced in stromal cells isolated and cultured from *Col5a1^d/d^* mice compared to those from *Col5a1^f/f^* mice, confirming efficient deletion. Similar to decidual tissues, *Prl8a2* gene expression was significantly reduced in *Col5a1^d/d^* cells compared to *Col5a1^f/f^* cells. In contrast to the findings in decidual tissues, *Gja1* (encoding connexin 43 gap junction protein) expression was significantly reduced in *Col5a1^d/d^* cells compared to *Col5a1^f/f^* cells in vitro (Fig. 4F). Collectively, these results demonstrate that type V collagen regulates a specific subset of decidualization pathways during embryo implantation.

To identify transcriptional pathways affected by deletion of *Col5a1* in the mouse decidua, we performed bulk RNA sequencing using total RNA isolated from GD6 decidual tissues of *Col5a1^f/f^* and *Col5a1^d/d^* mice. Principal component analysis (PCA) revealed group-specific clustering of samples (Fig. 5A). Quality-control metrics collected by the nf-core/rnaseq pipeline, including total number of reads and percentage of uniquely mapped reads, were assessed (Supplementary Fig. 6). A total of 464 genes were significantly downregulated, 424 genes were significantly upregulated, and 16,991 genes were not altered, as represented in a volcano plot (Fig. 5B). The 50 most differentially expressed genes—comprising the top upregulated and downregulated transcripts—were visualized in a hierarchically clustered heatmap (Fig. 5C). The top 20 transcription factors significantly altered in *Col5a1*-deficient uteri are depicted in Fig. 5D. To characterize the biological significance of these transcriptional changes, functional enrichment analysis was performed on significantly differentially expressed gene sets using Over-Representation Analysis (ORA) against Gene Ontology Biological Process (GO-BP) terms. Several GO-BP, KEGG, and Reactome clusters were significantly downregulated in *Col5a1^d/d^* decidua. Conversely, upregulated GO-BP, KEGG, and Reactome clusters in *Col5a1^d/d^* decidua included extracellular matrix organization, ECM–receptor interaction, focal adhesion, collagen degradation, and formation of fibrin clot (clotting cascade) (Fig. 5E). Gene Set Enrichment Analysis (GSEA) using Hallmark gene sets further identified pathways significantly altered in *Col5a1^d/d^* mice. Among the upregulated pathways were epithelial–mesenchymal transition, apical junction, TGF-β signaling, coagulation, and hypoxia, whereas downregulated pathways included DNA repair, fatty acid metabolism, Hedgehog signaling, interferon-gamma and interferon-alpha responses, and E2F targets (Fig. 5F).

Genes known to be involved in the decidualization process, such as *Fkbp5* and *Klf15*, were significantly reduced in *Col5a1^d/d^* decidua. Notably, in addition to *Prl8a2*, another prolactin family member, *Prl3c1* (encoding placental prolactin-like protein J [PLP-J]), was also significantly reduced (Fig. 6A). Both genes were induced in stromal cells during decidualization^28,29^. Other prolactin family members, such as *Prl3d1/Prl3d2* (encoding placental lactogen 1), *Prl5a1* (encoding placental prolactin-like protein L), *Prl7a1* (encoding prolactin-like protein E), and *Prl7a2* (encoding prolactin-like protein F), were significantly reduced in *Col5a1^d/d^* decidua (Fig. 6A). These genes are primarily expressed by trophoblast cells^30^. Despite the removal of embryos from the decidua during tissue collection prior to RNA extraction, the expression of these genes indicates that their transcripts were likely derived from invasive trophoblasts within the decidua. The reduced expression of these genes in *Col5a1^d/d^* decidua suggests that trophoblast invasion is compromised compared to *Col5a1^f/f^* decidua. The most significantly downregulated individual genes included *Adcy5*, *Fam20c*, *Bmper*, *Igfbp2*, and *Ccdc141*, whereas the most significantly upregulated genes included *Ltbp1*, *Igfbp5*, *Tnc*, *Slc16a2*, and *C1qtnf7*, as identified by this RNA-seq analysis (Fig. 6B). These findings suggest that deletion of Col5a1 selectively impairs pathways of endometrial decidualization in mice.

**Fig. 6.**
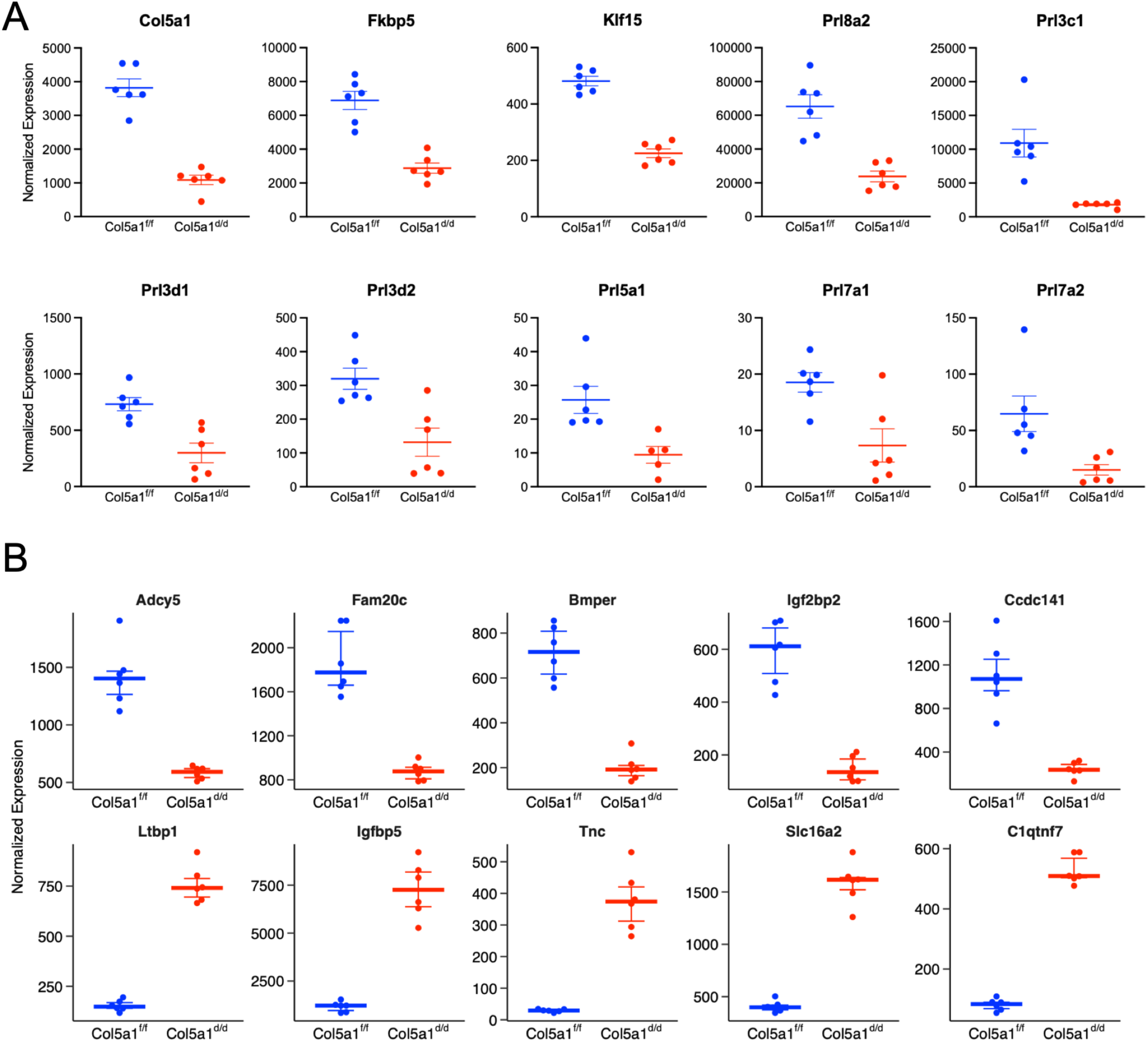
Selective genes differentially expressed in *Col5a1*-deficient uteri identified through RNA sequencing analysis. (A) Normalized expression of *Col5a1*, *Fkbp5*, *Klf15*, *Prl8a2*, *Prl3c1*, *Prl3d1*, *Prl3d2*, *Prl5a1*, *Prl7a1*, and *Prl7a2* between *Col5a1^f/f^*and *Col5a1^d/d^* decidual samples. (B) Top genes differentially expressed between *Col5a1^f/f^* and *Col5a1^d/d^* decidual samples. Genes shown meet both thresholds (|log₂ fold change| > 1 and adjusted *P* < 0.05).

### Type V collagen is crucial for reorganizing and maintaining the structural integrity of fibrillar collagen in the decidua

Given the notable enrichment of differentially expressed ECM-associated genes in the RNA-seq dataset, a focused analysis was conducted using the 580 differentially expressed genes classified against the Molecular Signatures Database (MSigDB) matrisome gene sets. This analysis identified 64 upregulated and 30 downregulated ECM-associated genes in *Col5a1^d/d^* decidua, as visualized in a volcano plot (Fig. 7A). Differential expression of ECM gene components was further stratified by matrisome subtypes and represented in a heatmap (Fig. 7B). Within the collagen subgroup (33 genes), *Col5a1* was expectedly downregulated in *Col5a1^d/d^* mice, whereas six collagen genes were upregulated, including *Col6a1*, *Col6a2*, *Col6a4*, *Col6a6*, *Col25a1*, and *Col26a1*. In the glycoprotein subgroup (126 genes), 16 genes were upregulated—including *Igfbp2*, *Igfbp5*, *Mfap4*, *Thbs1*, and *Fbln5*—while five genes were downregulated, including *Fgg*, *Eln*, *Thsd4*, *Gldn*, and *Bmper*. Within the proteoglycan subgroup (22 genes), all four differentially expressed genes were upregulated, including *Omd*, *Lum*, *Ogn*, and *Hapln3*. In the ECM-regulators subgroup (133 genes), 13 genes were upregulated— including *Loxl1*, *Loxl2*, *Adamts19*, *Mmp7*, and *Mmp12*—while seven were downregulated, including *Mmp1a*, *Adam11*, *Adamts18*, and *Fam20c*. Within the ECM-affiliated subgroup (91 genes), seven genes were upregulated, including *C1qtnf2*, *C1qtnf7*, *Clec21*, *Muc4*, *Muc20*, *Sftpd*, and *Sema3a*, whereas two genes were downregulated, *Lgals4* and *Muc13*. Finally, within the secreted factor subgroup (175 genes), 18 genes were upregulated—including *Cxcl5*, *Wnt9a*, and *Bmp7*—while 15 genes were downregulated, including *Wnt1*, *Bmp8a*, *Prl5a1*, and *Prl7a2*.

**Fig. 7.**
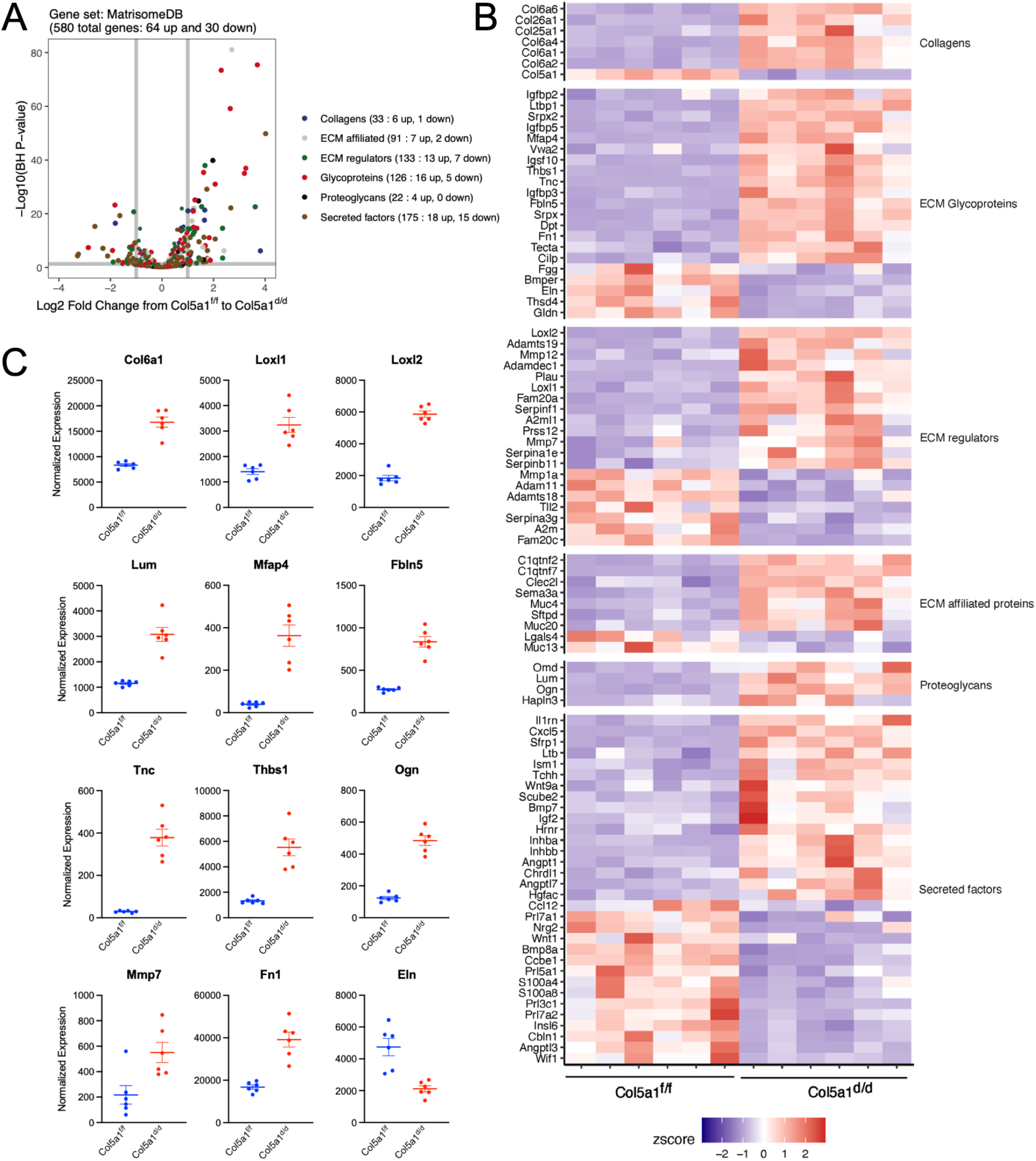
Deletion of *Col5a1* significantly impacts matrisome gene expression. (A) Volcano plot illustrating differential matrisome gene expression, with genes color-coded by matrisome category. (B) Heatmap illustrating differential expression of matrisome genes grouped by category. (C) Normalized expression of selected matrisome genes between *Col5a1^f/f^* and *Col5a1^d/d^* decidual samples. Genes shown meet both thresholds (|log₂ fold change| > 1 and adjusted *P* < 0.05).

Multiple genes encoding factors involved in the synthesis, processing, and assembly of fibrillar collagen were differentially expressed in *Col5a1^d/d^* decidua (Fig. 7C). However, many genes encoding factors involved in the synthesis, processing, and assembly of fibrillar collagen remained unchanged in *Col5a1^d/d^* decidua (Supplementary Fig. 7). The expression of *Col1a1* and *Col3a1*, genes predominantly encoding fibrillar collagen, was unchanged in *Col5a1^d/d^* decidua (Supplementary Fig. 7). Consistent with the gene expression data, the protein levels of COL1A1 remained unchanged between *Col5a1^f/f^*and *Col5a1^d/d^* decidua (Supplementary Fig. 8). These data suggest that other fibrillar collagens do not compensate for the loss of *Col5a1*. To evaluate fibrillar collagen reorganization, we localized COL1A1 protein in *Col5a1^d/d^* decidua. The expression pattern of COL1A1 was restricted to the immediate vicinity of the embryo in *Col5a1^d/d^* decidua, contrasting with *Col5a1^f/f^*decidua, where COL1A1 was extensively distributed throughout the decidua in an organized network (Fig. 8A).

**Fig. 8.**
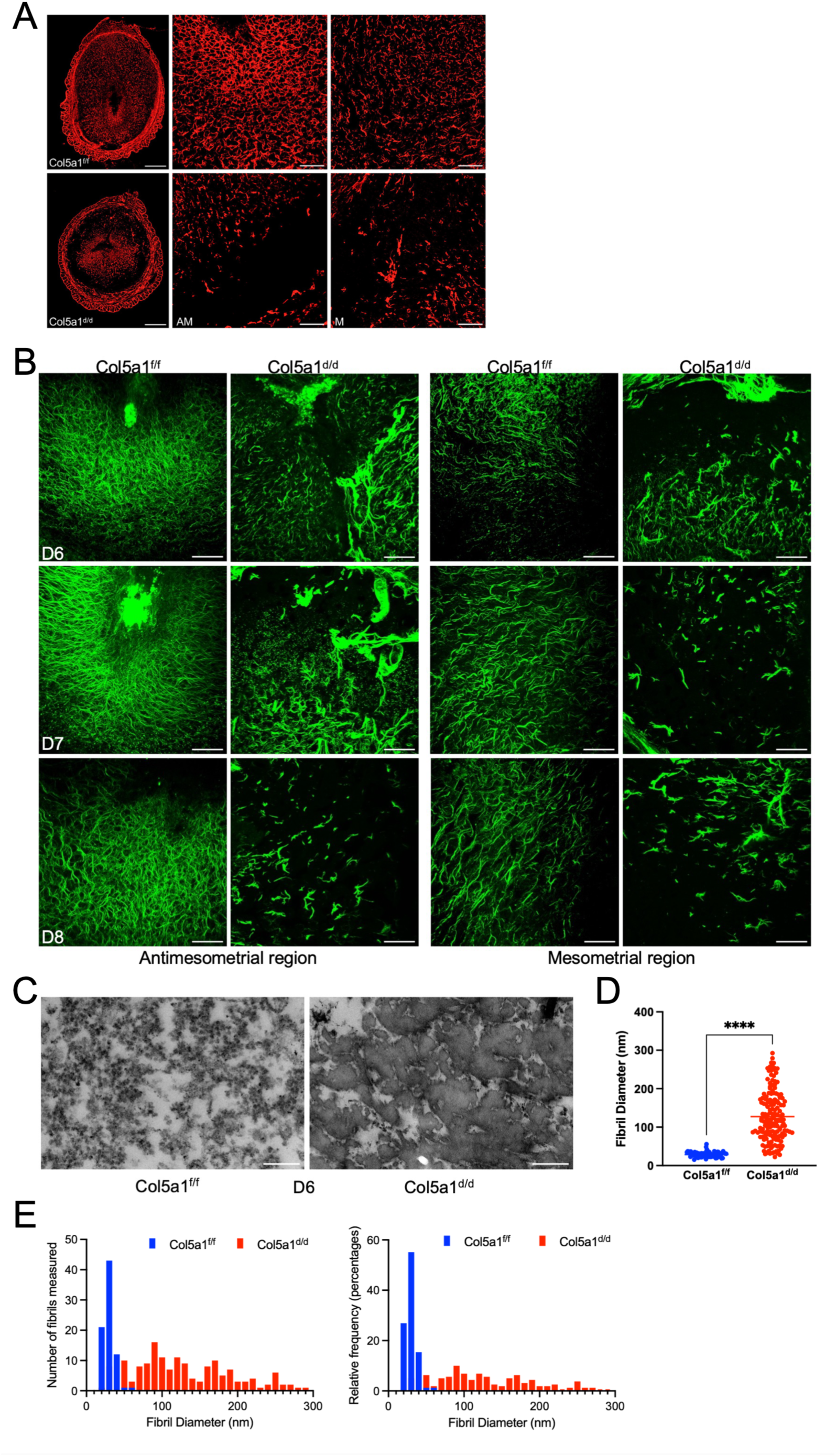
Disruption of fibrillar collagen reorganization in *Col5a1^d/d^* decidua. (A) Confocal imaging of COL1A1 in frozen uterine sections from *Col5a1^f/f^* and *Col5a1^d/d^* mice on GD6. AM, antimesometrial region; M, mesometrial region (n = 3 mice per genotype). Scale bar, 500 µm (left panel) and 100 µm (middle and right panels). (B) Second harmonic generation imaging of fibrillar collagen in frozen uterine sections from *Col5a1^f/f^*and *Col5a1^d/d^* mice from GD6–GD8 implantation sites. Images were captured in mesometrial and antimesometrial regions (n = 3 mice per time point per genotype). Image acquisition settings were optimized for morphology visualization. Scale bar, 100 µm. (C) Transmission electron microscopy images showing ultrastructural features of collagen fibrils in decellularized GD6 decidua from *Col5a1^f/f^* and *Col5a1^d/d^* uteri. Scale bar, 200 nm. (D) Quantification of collagen fibril diameter from decellularized GD6 decidua (n = 3 mice per genotype). A total of 78 (*Col5a1^f/f^*) and 161 (*Col5a1^d/d^*) fibrils were analyzed (****P < 0.0001). (E) Graphs showing the distribution and relative frequency of collagen fibril diameters.

To further evaluate fibrillar collagen reorganization within deeper regions of the decidua, we performed second harmonic generation imaging^13^. In *Col5a1^f/f^* decidua from GD6 through GD8, fibrillar collagen was extensively remodeled at the embryo implantation sites. Collagen fibers were dense and oriented around the embryo along the mesometrial–antimesometrial axis. In contrast, in *Col5a1^d/d^* decidua, the collagen fiber network was severely disrupted and collapsed (Fig. 8B). In the absence of type V collagen, fibrillar collagen failed to assemble into an organized network, as observed in *Col5a1^f/f^* decidua. To further examine fibrillar collagen defects, we evaluated ultrastructural features using transmission electron microscopy. In *Col5a1^f/f^*decidua, collagen fibrils were circular and uniform in size and shape. In contrast, in *Col5a1^d/d^* decidua, collagen fibrils were enlarged, irregular, and exhibited a cauliflower-like morphology, indicative of disrupted collagen fibrillogenesis (Fig. 8C). Fibril diameter was significantly greater in *Col5a1^d/d^* decidua compared to *Col5a1^f/f^* decidua (Fig. 8D). The relative frequency distribution of collagen fibril diameters between *Col5a1^f/f^* and *Col5a1^d/d^* decidua further reflected differences in fibril size distribution consistent with their altered morphology and impaired assembly (Fig. 8E). These findings demonstrate that type V collagen is necessary for fibrillar collagen processing, assembly, and reorganization during endometrial decidualization.

### Type V collagen regulates decidual angiogenesis and vascular network formation

Type V collagen is expressed in endothelial cells and in the vascular ECM of many tissues and has been demonstrated to be involved in angiogenesis. Therefore, we co-localized type V collagen with cluster of differentiation 31 (CD31), a marker of endothelial cells, to examine its expression in the decidual vasculature. On GD6, the vascular network within the decidua was less developed, CD31 staining was scarcely distributed throughout the decidua, and type V collagen showed limited alignment with CD31 (Fig. 9A). However, the vascular network was well developed by GD8, with extensive branching within the mesometrial decidua. At GD8, type V collagen exhibited a strong co-localization with CD31 throughout the decidua. Notably, this co-localization was particularly intimate with the vascular network in the mesometrial region (Fig. 9B). These findings suggest a role for type V collagen in angiogenesis, as development of the vascular network and its branching are more prominent in the mesometrial region, a presumptive site of placentation. To further evaluate the impact of *Col5a1* deletion on decidual angiogenesis and vascular network development, we localized CD31 within the decidua of *Col5a1^f/f^* and *Col5a1^d/d^* uteri. On GD6 and GD7, the vascular network formed appropriately in *Col5a1^f/f^*decidua, whereas the vascular network surrounding the implanting embryo was disrupted in *Col5a1^d/d^* decidua (Fig. 9C). By GD8, the characteristic and extensive branching vascular network was absent in the mesometrial region of *Col5a1^d/d^* decidua compared to *Col5a1^f/f^* decidua. In the antimesometrial region, vascular network density was markedly reduced in *Col5a1^d/d^* decidua compared to *Col5a1^f/f^* decidua (Fig. 9D).

**Fig. 9.**
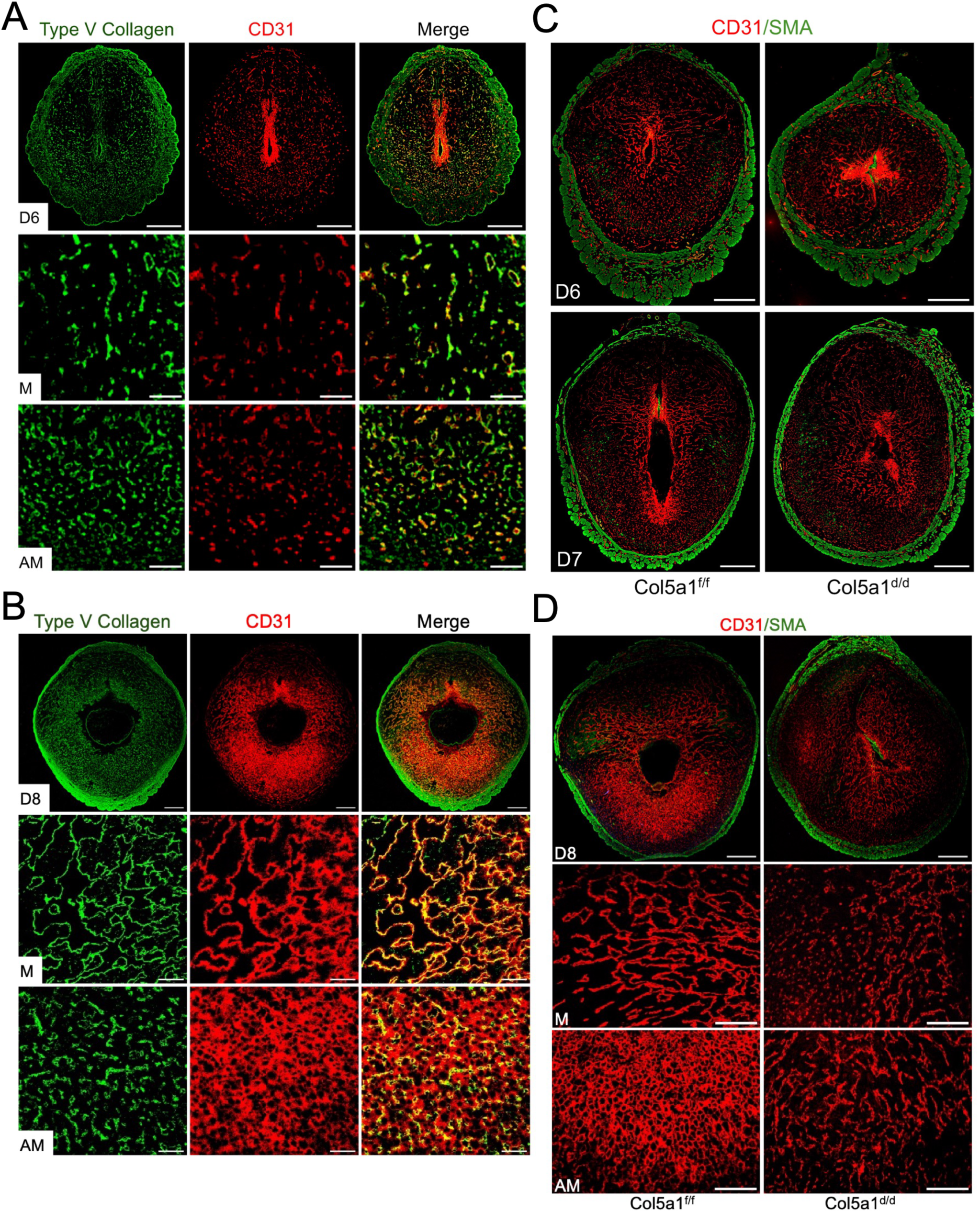
Deletion of COL5A1 impairs decidual angiogenesis in the mouse uterus. (A) Confocal microscopy images showing co-localization of COL5A1 (green) and CD31 (red) in GD6 uterine sections (n = 6 mice). Scale bar, 500 µm (top panel) and 100 µm (middle and lower panels). (B) Confocal microscopy images showing co-localization of COL5A1 (green) and CD31 (red) in GD8 uterine sections (n = 6 mice). Scale bar, 500 µm (top panel) and 100 µm (middle and lower panels). (C) Confocal microscopy images showing co-localization of CD31 (red) and smooth muscle actin (SMA; green) in GD6 and GD7 uterine sections from *Col5a1^f/f^* and *Col5a1^d/d^*mice (n = 3–4 mice per genotype). Scale bar, 500 µm. (D) Confocal microscopy images showing co-localization of CD31 (red) and smooth muscle actin (SMA; green) in GD8 uterine sections from *Col5a1^f/f^* and *Col5a1^d/d^* mice (n = 3–6 mice per genotype). Scale bar, 500 µm (top panel) and 100 µm (middle and lower panels). Image acquisition settings were optimized for morphology visualization. AM, antimesometrial region; M, mesometrial region.

Consistent with the gross morphological and histopathological evidence of intrauterine hemorrhage, and with the aberrant CD31 immunostaining pattern indicative of disrupted decidual vasculature in *Col5a1^d/d^* mice, we examined angiogenesis-associated gene sets within the differentially expressed transcriptome. We identified 110 differentially expressed genes out of a total of 528 genes, including 28 upregulated and 12 downregulated genes, as shown in a volcano plot (Fig. 10A). Hierarchical clustering of differentially expressed angiogenesis-related genes is presented in a heatmap (Fig. 10B). Among the downregulated genes were *Bmper*, *Angptl3*, *Sphk1, Hif3a*, *Ccbe1, Minar1,* and *Adra2b* (Fig. 10C). Angiopoietins and angiopoietin-like proteins were significantly dysregulated in the decidua lacking *Col5a1*. Specifically, *Angptl3* was significantly reduced, whereas *Angpt1*and *Angptl7* were increased (Fig. 10C, D). In contrast, *Angpt2, Angpt4, Angptl2, Angptl4,* and *Angptl6* remained unchanged (Supplementary Fig. 9). Furthermore, *Edn*, *Edn3*, and *Ednra*—genes encoding endothelins and their receptor—were abnormally elevated in *Col5a1^d/d^* decidua (Fig. 10D). Genes encoding key factors known to be involved in decidual angiogenesis, including vascular endothelial growth factor (VEGF) and hypoxia-inducible factors (HIFs), were not altered in *Col5a1*-deficient decidua (Supplementary Fig. 9). Consistently, the localization pattern of VEGF was similar between *Col5a1^f/f^* and *Col5a1^d/d^* decidua (Fig. 10E). Expression levels of *Hif1a* and *Epas1* (*Hif2a*), which are known regulators of decidualization, were also unchanged (Supplementary Fig. 9). However, the spatial distribution of HIF2A differed, with localization restricted to decidual cells surrounding the embryo in *Col5a1^d/d^* tissue, compared to a broader distribution in *Col5a1^f/f^* decidua (Fig. 10F). Collectively, these findings indicate that loss of type V collagen severely impairs decidual angiogenesis and disrupts the development of the decidual vascular network.

**Fig. 10.**
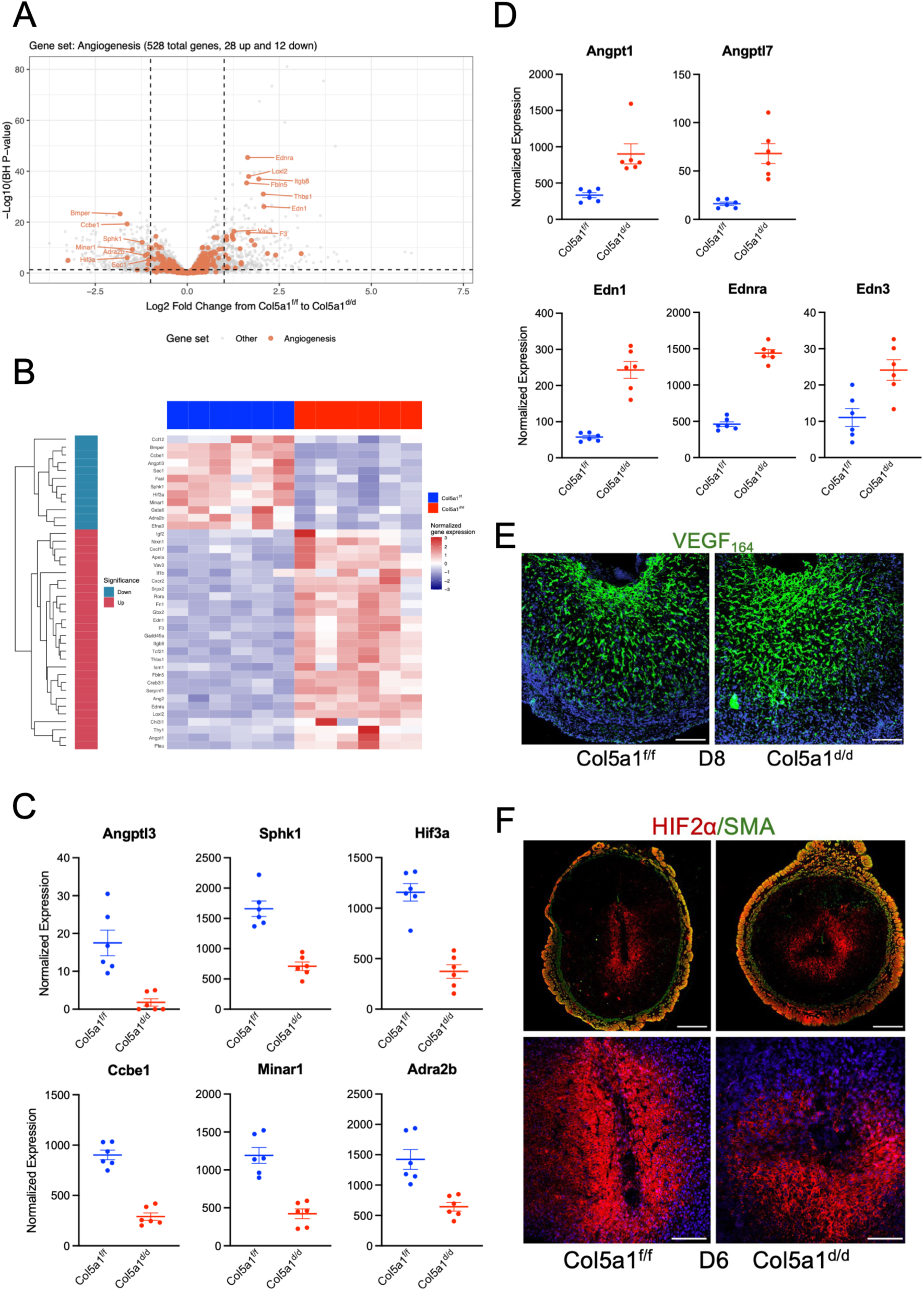
Loss of type V collagen impairs decidual angiogenesis. (A) Volcano plot of differential expression of angiogenesis-related genes showing genes with large expression changes and strong statistical significance located furthest from the center and highest on the plot. (B) Heatmap showing the top 40 differentially expressed angiogenesis-related genes ranked by adjusted *P* < 0.05. Blue indicates downregulation and red indicates upregulation in *Col5a1^d/d^*decidua. (C) Normalized expression of angiogenesis-related genes downregulated in *Col5a1^d/d^* decidua identified by RNA sequencing. Genes shown meet both thresholds (|log₂ fold change| > 1 and adjusted *P* < 0.05). (D) Normalized expression of angiogenesis-related genes upregulated in *Col5a1^d/d^* decidua identified by RNA sequencing. Genes shown meet both thresholds (|log₂ fold change| > 1 and adjusted *P* < 0.05). (E) Confocal microscopy images showing localization of VEGF164 (green) in GD8 uterine sections from *Col5a1^f/f^* and *Col5a1^d/d^* mice. DAPI (blue) (n = 3–4 mice per genotype). Scale bar, 200 µm. (F) Confocal microscopy images showing localization of HIF2α (red) in GD6 uterine sections from *Col5a1^f/f^* and *Col5a1^d/d^* mice. Image acquisition settings were optimized for morphology visualization. SMA (green); DAPI (blue) (n = 3–5 mice per genotype). Scale bar, 500 µm (top panel) and 100 µm (lower panel).

## DISCUSSION

This study uncovers a previously unrecognized role of type V collagen in ECM reorganization and angiogenesis during endometrial decidualization at the time of embryo implantation, thereby contributing to the establishment of the maternal–fetal interface required for a successful pregnancy. Using a uterine- specific *Col5a1* conditional knockout mouse model, we demonstrate that loss of type V collagen results in complete pregnancy loss due to severe structural distortion of the decidua, profound disorganization of fibrillar collagen, shallow and misdirected embryo invasion, and disruption of the decidual vascular network. COL5A1 is essential for type V collagen synthesis and serves as an obligatory chain in the formation of heterotrimeric collagen molecules^19^. Type V collagen, although a minor fibrillar collagen, is a critical regulator of fibrillar collagen synthesis and assembly^20^. Embryonic lethality observed in *Col5a1* mutant mouse models, together with Ehlers–Danlos syndrome caused by COL5A1 mutations in humans, highlights the fundamental importance of type V collagen in maintaining ECM structure and normal tissue function^20,22^. The indispensable role of type V collagen in fibrillar collagen assembly and ECM organization, and thereby in maintaining the integrity and function of collagen-rich tissues such as tendon, cornea, and skin, has been extensively documented^22^. In addition to connective tissues, the functional importance of type V collagen in soft and vital tissues such as the heart and kidney has also been reported^31,32^. In this study, we extend these findings to another soft tissue, the uterus, demonstrating a critical role for type V collagen in supporting reproductive function and establishing a successful pregnancy.

The endometrium forms a transient tissue structure known as the decidua to support embryo implantation. We previously reported that fibrillar collagens are extensively synthesized, reorganized, and aligned parallel to the direction of embryo invasion within the decidua^13^. In this study, we provide compelling evidence that fibrillar collagen reorganization is crucial for embryo implantation and for establishing the maternal–fetal interface during early pregnancy. Embryo attachment and invasion trigger extensive endometrial stromal cell proliferation and differentiation, resulting in decidua formation. Although deletion of *Col5a1* did not affect stromal cell proliferation per se, it significantly affected specific differentiation pathways. One of the notably affected pathways is the prolactin gene family. In particular, *Prl8a2* (prolactin family 8, subfamily a, member 2), also known as decidual prolactin-related protein (dPRP), a key signaling hormone secreted by the mouse uterine decidua, was significantly downregulated in the absence of *Col5a1*. These findings indicate a selective impairment of the decidualization process. RNA-sequencing analysis revealed several molecular pathways potentially responsible for impaired decidualization, including alterations in ECM reorganization and disrupted angiogenesis and vascular network formation in the uterus lacking *Col5a1*.

Fibrillar collagens, the predominant fibrous components of the ECM, are ubiquitously expressed in almost all tissues of the body^16^. The structure and composition of the ECM are established during embryonic development and undergo continuous but controlled remodeling during adult life to maintain tissue mechanical homeostasis^15,33^. In contrast to ECM-rich connective tissues, reproductive tissues such as the uterus undergo repeated cycles of extensive growth, remodeling, and involution to support physiological functions during pregnancy and parturition^34–36^. Tissues with abnormal fibrillar collagen structure exhibit loss of integrity and function, as has been reported in multiple collagen mutant mouse models and human genetic conditions^18,31,36^. In this study, we disrupted fibrillar collagen structure through deletion of *Col5a1* and evaluated its functional importance during embryo implantation. Loss of *Col5a1* resulted in abnormal collagen fibril ultrastructure due to impaired fibrillogenesis. Microscopic analyses further demonstrated that collagen fibers were disorganized within the decidua. Consistent with these structural abnormalities, the gross morphology of the decidua was severely distorted, indicating that proper fibrillar collagen reorganization is required to maintain decidual integrity for embryo invasion. Due to loss of fibrillar collagen orientation and compromised decidual tissue integrity, embryo invasion was disrupted within the decidua. Although embryos were able to breach the luminal epithelium and enter the uterine stroma, their subsequent invasion and embedding were misdirected. Furthermore, invading embryos failed to establish stable interactions with the surrounding decidual tissue, which are necessary for continued survival and development. These findings provide insight into the long-standing question of how embryo invasion is directed along the mesometrial–antimesometrial axis within the decidua. They also offer mechanistic clues regarding the migration of invasive trophoblast cells within this tissue. It is well established that tumor cells preferentially migrate along collagen fibers within the primary tumor microenvironment, a process facilitated by collagen organization^37,38^. Although real-time intravital imaging at the maternal–fetal interface remains technically challenging, development of alternative approaches to study fibrillar collagen-guided trophoblast migration will be important for future investigations.

Decidual angiogenesis involves the formation of an extensive vascular network within the rapidly developing and transient decidual tissue^11^. This vascular network is essential not only for maintaining decidual structure but also for supporting and nourishing the invading embryo until placental development is complete. Multiple decidual stromal-derived factors have been implicated in this process^11,39,40^. In this study, we identify a previously unrecognized role for type V collagen in regulating decidual angiogenesis. Type V collagen is abundantly expressed in decidual endothelial cells. Deletion of *Col5a1* resulted in catastrophic uterine hemorrhage, followed by complete embryo resorption, due to severe disruption of the vascular network and failure of angiogenic processes within the decidua. This angiogenic failure may arise from two potential mechanisms. First, distorted decidual structure caused by defective ECM assembly may fail to provide a stable scaffold to support vessel sprouting and may disrupt the proper distribution of angiogenic factors. Second, intrinsic loss of type V collagen in endothelial cells may impair their ability to support vessel formation and compromise vascular integrity. These findings reveal a previously unappreciated role of type V collagen beyond its canonical function in fibrillar collagen assembly, identifying it as a critical regulator of angiogenesis during embryo implantation.

Multiple angiogenesis-related pathways are altered in the *Col5a1^d/d^*decidua. Angiopoietins and angiopoietin-like proteins are highly expressed in the mouse decidua^39,40^, and several members of this family are differentially regulated in *Col5a1^d/d^* tissue. While *Angpt1* and *Angptl3* function as pro-angiogenic molecules, *Angptl7* acts as an anti-angiogenic factor and has been implicated in ECM reorganization^41^. In our study, deletion of Col5a1 resulted in aberrant activation of endothelin signaling. Endothelins and their receptors are expressed in the human decidua^42^. Endothelin-1 has been shown to inhibit decidualization of human endometrial stromal cells in vitro^43^. In addition, endothelins are recognized for their roles in regulating collagen synthesis and ECM remodeling across multiple tissues^44,45^. Elevated maternal expression of Edn1 has been associated with impaired implantation in mice, characterized by reduced vascular density in the mesometrial region of the decidua and abnormal ectoplacental cone morphology, independent of embryo genotype^43,46^. Notably, these detrimental effects were mitigated by nicotinamide supplementation^46^. Female mice deficient in Sphk1 and Sphk2 (Sphk1^−/−^Sphk2^+/−^) exhibit impaired decidualization, marked by severely compromised uterine vasculature and resulting in early pregnancy loss^47^. Moreover, Sphk1 expression is influenced by ECM stiffness^48^. Although the interactions among these pathways in the decidua require further investigation, their dysregulation may collectively contribute to the severe phenotype observed in *Col5a1^d/d^*mice.

Our data suggest a mechanistic framework linking defects in fibrillar collagen reorganization to impaired endometrial decidualization and pregnancy failure. These findings may have relevance to human reproductive disorders associated with defective decidualization and vascular remodeling, including recurrent pregnancy loss, implantation failure, fetal growth restriction, and preeclampsia^49–51^. This study also provides insight into clinical complications such as shallow implantation and placenta accreta spectrum disorders^51,52^. ECM reorganization during decidualization may play a crucial role in precisely regulating the depth and direction of embryo invasion. Defective ECM reorganization could lead to excessive trophoblast invasion into the myometrium, resulting in placenta accreta, whereas insufficient invasion may lead to shallow implantation^53^. An alternative concept suggests that disrupted collagen architecture at prior uterine scar sites is a defining feature of abnormal placental adherence in placenta accreta spectrum disorders^54^. Mutations in *COL5A1* are the primary cause of classical Ehlers–Danlos syndrome^21^. Women with EDS exhibit an increased incidence of infertility, spontaneous abortions, preterm labor, abnormal uterine bleeding, dysmenorrhea, and severe dyspareunia^55^. Our findings provide mechanistic insight into how loss of COL5A1 function in the uterus may contribute to pregnancy failure. Structural alterations in the ECM and the associated decline in tissue function are hallmarks of aging and are widely observed across connective tissues^56^. Although the impact of reproductive aging on fibrillar collagen structure—and its contribution to reproductive decline—remains poorly understood, this relationship warrants further investigation, particularly in the context of the increasing average maternal age at childbirth. Notably, the incidence of serious pregnancy complications, including miscarriage, rises significantly with advancing maternal age^57^.

In summary, conditional deletion of *Col5a1* in the decidua identifies type V collagen as a pivotal regulator of ECM architecture and angiogenic signaling during early pregnancy. By establishing a link between ECM defects and impaired decidual function leading to pregnancy loss, this study advances our understanding of how the maternal uterine environment is organized to support formation of the maternal– fetal interface. Furthermore, these findings highlight the potential of ECM-targeted pathways as diagnostic and therapeutic targets for uterine causes of reproductive failure.

## MATERIALS AND METHODS

### Animals

All animal procedures were performed in accordance with the National Institutes of Health Guide for the Care and Use of Laboratory Animals and were approved by the Institutional Animal Care and Use Committee at the University of Vermont. Mice harboring loxP sites flanking exons 3 and 4 of *Col5a1* (*Col5a1^f/f^*) were crossed with mice expressing Cre recombinase from a single allele of the progesterone receptor locus (*Pgr^cre/+^*)^18,58^. This breeding strategy generated *Pgr^cre/+^Col5a1^f/f^* females (referred to as *Col5a1^d/d^*). Littermate or age matched *Col5a1^f/f^* and *Col5a1^d/d^* mice were used in all experiments. Female *Col5a1^f/f^* and *Col5a1^d/d^* mice (2 to 3 months old) were mated with male mice as individual pairs and monitored for litter size and the number of pups born over a period of six months to assess their reproductive phenotype.

### Tissue collection

To collect uteri at specific time points, *Col5a1^f/f^* and *Col5a1^d/d^* female mice were mated with *Col5a1^f/f^* male mice for 6–8 h during the daytime. The day of the vaginal plug was considered day 0 of pregnancy (GD0). Reproductive tissues were collected between GD3 and GD12. Uteri were dissected, trimmed of adipose tissue, rinsed in PBS, and implantation sites were collected. Tissues were fixed in 10% neutral buffered formalin and paraffin embedded for histological analysis. For confocal imaging, tissues were freshly embedded in OCT compound (Tissue-Tek, Thermo Fisher Scientific, Waltham, MA, USA). For RNA and protein analysis, tissues were snap-frozen in liquid nitrogen and stored at −80 °C until further processing.

### Histopathology

Tissues were fixed in 10% neutral buffered formalin and paraffin embedded. Sections (5 µm) were cut on a microtome, air-dried at room temperature, and baked at 60 °C for 1 h. Slides were deparaffinized in xylene (three times) and rehydrated through a graded ethanol series (100%, 95%, 70%, and 50%; 5 min each), followed by rinsing in tap water for 10 min. Hematoxylin and eosin staining was performed using an H&E Stain Kit (Cat. No. H-3502, Vector Laboratories). Sections were stained with hematoxylin for 5 min and washed twice in distilled water for 15 s each. To neutralize residual acidity and enhance nuclear blue staining, slides were incubated in bluing reagent for 10 s and washed twice again in distilled water for 15 s each. Slides were then briefly immersed in 100% ethanol for 10 s, stained with eosin Y for 2–3 min, and rinsed in 100% ethanol for 10 s. Dehydration was completed in three changes of 100% ethanol for 1–2 min each. Slides were mounted with VectaMount Express Mounting Medium (Cat. No. H-5700- 60, Vector Laboratories). Whole uterine sections were captured using the Leica VERSA8 whole slide imager and subsequently processed using Aperio ImageScope and ImageJ software.

### Immunohistochemistry

Paraffin-embedded embryo implantation sites were sectioned at 5 µm. Slides were dried, heated at 60 °C for 1 h, deparaffinized, and rehydrated, followed by rinsing in tap water for 10 min. Antigen retrieval was performed in 10 mM sodium citrate buffer by microwave heating for 25 min. Endogenous peroxidase activity was quenched with 1% hydrogen peroxide in methanol for 15 min. Slides were washed in TBST for 5 min and blocked with normal goat serum (Cat. No. 50062Z, Thermo Fisher Scientific) in a humidified chamber for 30 min at room temperature. Sections were incubated overnight at 4 °C with primary antibodies against progesterone receptor (PGR; 1:200, rabbit, Cat. No. MA5-14505, Invitrogen), estrogen receptor alpha (ESR1; 1:100, Cat. No. sc-8005, Santa Cruz Biotechnology), or Ki-67 (MKI67; 1:100, Cat. No. 55609, BD Pharmingen). On the following day, slides were washed three times in TBST for 5 min each and incubated for 1 h at room temperature in a humidified chamber with the appropriate biotinylated secondary antibody: anti-mouse IgG (H+L) (Cat. No. BP-9200, Vector Laboratories) or anti-rabbit IgG (H+L) (Cat. No. BP-9100, Vector Laboratories), according to the host species of the primary antibody. After three additional washes in TBST, slides were incubated with streptavidin–peroxidase (ready-to-use; Cat. No. SA-5704, Vector Laboratories) for 30 min at room temperature and developed with DAB substrate (Cat. No. SK-4105, Vector Laboratories). Sections were then blued in 1.5% NH₄OH in 70% ethanol for 1 min, rinsed in NH₄OH in water, washed with tap water, and mounted with VectaMount Express Mounting Medium (Cat. No. H-5700-60, Vector Laboratories). Whole uterine sections were captured using the Leica VERSA8 whole slide imager and subsequently processed using Aperio ImageScope and ImageJ software.

### Confocal microscopic imaging

Frozen uterine cross-sections were fixed in acetone for 10 min and blocked with 10% normal goat serum (Life Technologies) for 30 min at room temperature. Primary antibodies were diluted in blocking solution, applied to tissue sections, and incubated overnight at 4 °C. Primary antibodies included collagen type I alpha 1 chain (COL1A1; 1:200, Cat. No. 72026, Cell Signaling Technology), collagen V (1:1000, Cat. No. ab7046, Abcam), pan-keratin (1:100, Cat. No. 4545, Cell Signaling Technology), trophoblast-specific protein alpha (1:250; TPBPA; Cat. No. ab104401, Abcam), placental lactogen 1 (1:250; PL-1; 1:100, Cat. No. sc-376436, Santa Cruz Biotechnology), CD31 (1:500, Cat. No. 553370, BD Pharmingen), desmin (1:250; Cat. No. 5332, Cell Signaling Technology), vascular endothelial growth factor A (1:10; VEGFA, Cat. No. AF-493-NA, R&D Systems), hypoxia-inducible factor 2 alpha (1:100; HIF2A; Cat. No. NB100- 122, Novus Biologicals), and alpha-smooth muscle actin (ACTA2; 1:1000, Cat. No. sc-32251, Santa Cruz Biotechnology). On the following day, sections were washed with PBS and incubated for 30 min at room temperature with species-specific secondary antibodies diluted in blocking solution. Secondary antibodies included Alexa Fluor 555–conjugated anti-rabbit IgG (1:500, Cat. No. A32732, Thermo Fisher Scientific) and Alexa Fluor 488–conjugated anti-mouse IgG (1:500, Cat. No. A11029, Thermo Fisher Scientific). Sections were washed with PBS and mounted with ProLong Gold Antifade Mountant containing DAPI (Thermo Fisher Scientific). A Nikon A1R confocal microscope (Nikon Instruments) was used to generate the images; this microscope is equipped with a galvanometer scanner and point-scan illumination at a rate of eight frames per second for a 1024 × 1024-pixel field. Both 4× and 10× objective lenses were used for imaging. Nikon NIS-Elements software (Nikon Instruments, Melville, NY) was used for image acquisition. ImageJ was used for image analysis.

### Second harmonic generation imaging

Frozen uterine cross-sections (50 µm) containing implantation sites from GD6–GD8 were thawed, covered with 0.1 M phosphate-buffered saline (PBS), and imaged using a Zeiss LSM 7 inverted microscope (Carl Zeiss AG, Oberkochen, Germany) equipped with an Achroplan 20×/0.8 objective lens. A Chameleon XR pulsed Ti:sapphire laser (Coherent, Santa Clara, CA, USA) tuned to 900 nm was used for excitation, and second harmonic generation signals were detected at 450 nm. Acquired images were analyzed using ImageJ.

### Alkaline phosphatase staining

Frozen sections of GD6 implantation sites were fixed in cold acetone for 10 min and washed three times in PBS for 5 min each. Slides were then incubated for 30 min in a humidified chamber with working solution prepared from an alkaline phosphatase detection kit according to the manufacturer’s instructions (Vector Laboratories, Blue Substrate Kit, Cat. No. SK-5300). Sections were washed with PBS followed by distilled water, air-dried for 1 h, and mounted with VectaMount. Whole uterine sections were captured using the Leica VERSA8 whole slide imager and subsequently processed using Aperio ImageScope and ImageJ software.

### Transmission electron microscopy

To decellularize implantation sites collected on GD6, tissues were incubated in 1% SDS in PBS with shaking for 2 h. Samples were then washed for 1 week in washing buffer, which was changed daily (0.9% NaCl, 0.05 M MgCl₂, 0.1 mg/mL DNase I, and 1% penicillin–streptomycin). Decellularized implantation sites were fixed overnight at 4 °C in half-strength Karnovsky’s fixative containing 2.5% glutaraldehyde, paraformaldehyde, and 0.05 M cacodylate buffer (pH 7.2). Samples were rinsed three times for 5 min each in cacodylate buffer, post-fixed in osmium tetroxide, and stained en bloc with 2% tannic acid in cacodylate buffer followed by 2% uranyl acetate. Tissues were dehydrated in graded ethanol, gradually infiltrated with propylene oxide:Spurr’s resin, embedded in fresh 100% Spurr’s resin, and polymerized at 70 °C. Ultrathin sections (80 nm) were cut using a Reichert Ultracut-II ultramicrotome with a Diatome Ultra 45 diamond knife, mounted on 200-mesh copper grids, stained with uranyl acetate and lead citrate, and examined using a JEOL 1400 transmission electron microscope (JEOL USA, Inc.) operated at 60 or 80 kV. Digital images were captured using an AMT-XR611 11-megapixel CCD camera (Advanced Microscopy Techniques). The diameter of each individual fibril was determined by drawing perpendicular lines to their longitudinal axis. Fibrils that overlapped or exhibited poor resolution were excluded from the measurements. Subsequently, diameters from multiple fibrils were aggregated to compute the mean diameter, count, and frequency distribution. Images were further processed and analyzed using ImageJ.

### Endometrial stromal cell culture

Endometrial stromal cells were isolated from GD3 uteri. Uteri were dissected, opened longitudinally, and cut into 3–5 pieces. Tissue digestion was performed in HBSS (Thermo Fisher Scientific) containing 6 g/L dispase (Cat. No. 07923, STEMCELL Technologies), 25 g/L pancreatin, and antibiotic–antimycotic solution (Thermo Fisher Scientific) for 1 h at room temperature followed by 15 min at 37 °C. Tissues were then digested in 2 mL HBSS containing 250 µg/mL Liberase (Sigma-Aldrich) at 37 °C for 45 min. Enzymatic digestion was terminated by the addition of 10% fetal bovine serum (Thermo Fisher Scientific). Stromal cells were collected by filtration through a 70-µm cell strainer followed by centrifugation at 300 × g for 5 min. Cell pellets were resuspended in DMEM/F12 (Thermo Fisher Scientific) supplemented with 2% FBS, 1% antibiotic–antimycotic solution, 10 nM 17β-estradiol (Sigma-Aldrich), and 1 µM progesterone (Sigma-Aldrich). Cells were plated in six-well plates at 1 × 10⁶ cells per well. Culture medium was changed after 2 h and subsequently every 24 h. Cell lysates were collected for gene and protein expression analyses.

### RNA isolation and qPCR

Total RNA was extracted using the RNeasy Mini Kit (Cat. No. 74104, Qiagen) according to the manufacturer’s protocol and reverse transcribed into cDNA using iScript Reverse Transcription Supermix (Bio-Rad Laboratories). Quantitative PCR was performed using SYBR Green master mix (Thermo Fisher Scientific). Gene-specific primers were designed for the genes of interest (Supplementary Table 3). Relative expression levels were normalized to the housekeeping gene *Rplp0* and calculated using the 2^−ΔΔCt^ method.

### Western blot

Tissues or cells were lysed in radioimmunoprecipitation assay (RIPA) buffer supplemented with 1% protease and phosphatase inhibitors (Thermo Fisher Scientific). Protein concentrations were determined using a bicinchoninic acid (BCA) assay according to the manufacturer’s instructions (Thermo Fisher Scientific). Protein samples were mixed with Laemmli sample buffer (Bio-Rad Laboratories) and β-mercaptoethanol (Sigma-Aldrich) and boiled at 95 °C for 10 min. Protein standards (Precision Plus Protein Kaleidoscope, Bio-Rad Laboratories) and samples were resolved on 10% Bis-Tris-HCl sodium dodecyl sulfate–polyacrylamide gel electrophoresis (SDS-PAGE) gels at 50 V for 10 min followed by 100 V for 1 h. Proteins were transferred onto nitrocellulose membranes (Bio-Rad Laboratories) at 100 V for 1 h at 4 °C. Membranes were blocked in 3% blotting-grade nonfat dry milk in TBST (Bio-Rad Laboratories) for 1 h at room temperature and incubated overnight at 4 °C with primary antibodies diluted in blocking solution. Primary antibodies included COL1A1 (1:1000, Cat no: 72026, Cell Signaling), collagen V (1:500; Catalog Number: 67604-1-Ig; Proteintech) and β-actin (1:1000, Catalog No. 3700S, Cell Signaling Technology). On the following day, membranes were incubated for 1 h at room temperature with HRP-conjugated secondary antibodies, goat anti-rabbit IgG (H+L) (1:1000, Cat. No. 7074S, Cell Signaling Technology) or goat anti-mouse IgG (H+L) (1:1000, Cat. No. 7076S, Cell Signaling Technology). Signal detection was performed using Amersham ECL Western Blotting Detection Reagents, and images were acquired using an ImageQuant 800 western blot imaging system (Cytiva Life Sciences).

### mRNA library preparation and sequencing

RNA-seq libraries were prepared using the KAPA/Roche mRNA HyperPrep Kit (KAPA/Roche, Cat. No. 08098123702) according to the manufacturer’s instructions. Briefly, 1 µg of total RNA was enriched using oligo(dT)-coupled magnetic beads. Purified mRNA was fragmented at 94 °C for 6 min and immediately placed on a magnet, and the supernatant was transferred to a new tube on ice. First-strand cDNA synthesis was followed by second-strand synthesis. Diluted KAPA unique dual indices (7 µM; KAPA/Roche, Cat. No. 08861919702) were ligated to the libraries, and products were purified using KAPA Pure Beads (KAPA/Roche, Cat. No. 07983298001). Libraries were then amplified, bead-purified, and eluted in 20 µL of 10 mM Tris-Cl, pH 8.5 (Qiagen, Cat. No. 19086). Library concentrations were determined using a Qubit 2.0 fluorometer (Thermo Fisher Scientific) with the dsDNA HS Assay Kit (Cat. No. Q32854), and library quality was assessed using a TapeStation 4200 system (Agilent, Cat. No. G2991BA) with High Sensitivity D5000 ScreenTape and associated reagents (Agilent, Cat. No. 5067-5592). Libraries were sequenced at the Hubbard Center for Genome Studies (University of New Hampshire) on a NovaSeq 6000 platform (Illumina) using paired-end version 1.5 chemistry on an SP patterned flow cell. Forward and reverse read lengths were 250 bp, and index reads consisted of dual 8-mers. Base calling and demultiplexing were performed using Illumina bcl2fastq v2.20.0.422.

### RNA-seq Data Processing

Raw paired-end RNA-seq reads were processed using the nf-core/rnaseq pipeline (version 3.25.0; commit 891468c) executed via Nextflow^59^ with Apptainer containerization. Briefly, reads were aligned to the mouse reference genome GRCm39 (Ensembl primary assembly) using STAR^60^. Genome annotation was obtained from GENCODE vM38 (basic annotation). Salmon^61^ was used to generate length-scaled TPM back-calculated counts. Differential expression analysis was performed in R (v 4.5.1) using DESeq2 (v 1.50.2)^62^ contrasting *Col5a1^d/d^* vs *Col5a1^f/f^*. Low-count genes were removed by retaining only those with ≥ 10 counts in at least three samples. The DESeq function was applied to estimate size factors, dispersion, and to fit negative binomial generalized linear models with Wald tests. Regularized log (rlog) transformation was applied for quality-control visualizations, including PCA and sample-to-sample distance clustering. Genes were considered significantly differentially expressed at an adjusted p-value (Benjamini–Hochberg FDR) < 0.05 and |log₂ fold change| ≥ 1.

Over-representation analysis (ORA) was performed on significantly up- and downregulated gene sets separately using *clusterProfiler*^63^ against Gene Ontology Biological Process (GO:BP), KEGG, and Reactome pathway databases. The full set of tested genes served as the statistical background. P-values were adjusted using the Benjamini–Hochberg method. Gene symbol-to-Entrez ID mapping was performed using *org.Mm.eg.db*. Gene Set Enrichment Analysis (GSEA) was performed using *clusterProfiler::GSEA* against the MSigDB Hallmark gene set collection for *Mus musculus*, retrieved via *msigdbr*. Genes were ranked based on the DESeq2 Wald statistic. P-values were adjusted using the Benjamini–Hochberg method. Transcription factor activity inference was performed using VIPER^64^ with regulons from the DoRothEA^65^ resource, restricted to confidence levels A, B, and C. A normalized enrichment score (NES) was computed for each transcription factor, with positive NES indicating inferred activation and negative NES indicating inferred repression in *Col5a1^d/d^* relative to *Col5a1^f/f^*. The angiogenesis gene set was downloaded from MSigDB^66^; mouse GOBP_ANGIOGENESIS (GO:0001525). Matrisome gene sets are all categories for mouse and downloaded from MatrisomeDB 2.0^67^ (https://matrisomedb.org/). All analyses were performed in R (version ≥ 4.3). Visualization was carried out using *ggplot2*, *pheatmap*, and *ggrepel*. The full analysis pipeline is reproducible via *run_all.R*.

### Statistics

Data were analyzed using GraphPad Prism software (GraphPad Software). Comparisons between two groups were performed using unpaired two-tailed Student’s *t*-tests. Comparisons among multiple groups were performed using one-way analysis of variance (ANOVA) followed by Tukey’s multiple comparisons test. Data are presented as mean ± SEM. A P value < 0.05 was considered statistically significant.

## Supporting information

Gebril et al 2026 Supplementary File

## Acknowledgements

Research reported in this publication was supported by the Eunice Kennedy Shriver National Institute of Child Health & Human Development of the National Institutes of Health under Award Numbers R01HD111479 (SN) and R00HD090301 (SN), and by the University of Vermont Cardiovascular Research Institute Early Career Research Award (SN). The content is solely the responsibility of the authors and does not necessarily represent the official views of the National Institutes of Health. We sincerely acknowledge Dr. David E. Birk, Ph.D., Distinguished Professor, Department of Molecular Pharmacology & Physiology, University of South Florida Morsani College of Medicine, for providing the *Col5a1* loxP mouse model, and Dr. Francesco DeMayo, Ph.D., Senior Investigator, Reproductive and Developmental Biology Laboratory / Pregnancy and Female Reproduction Group, NIEHS, for providing the *Pgr-cre* mice used in this study. We also thank Dr. Douglas Taatjes, Director, Nicole Bouffard, and Brad P. Vietje at the Microscopy Imaging Center at the University of Vermont (RRID: SCR_018821) for their help and support. Confocal microscopy was performed using a Nikon A1R-HD point-scanning confocal microscope supported by NIH award number 1S10OD025030-01 from the Office of Research Infrastructure Programs. We thank Dr. Todd Clason, Director of the Customized Physiology and Imaging Core at the University of Vermont, for his support in performing second harmonic generation imaging. Funding was also provided by P20 GM135007 from the National Institute of General Medical Sciences (NIGMS) of the NIH.

## Author contributions

SN conceptualized and designed the study. EK, MG, SN, and RD conducted experiments and analyzed the data. JB analyzed the RNA sequencing data and generated figures from this analysis. JPL provided the *Pgr-cre* mice used in this study. MG and SN wrote the manuscript. All authors reviewed and approved the final manuscript.

## Ethics declarations Conflict of interest

The authors declare no competing interests.

## Ethical approval

All animal experiments were conducted in accordance with the National Institutes of Health Guide for the Care and Use of Laboratory Animals and were approved by the Institutional Animal Care and Use Committee at the University of Vermont.

## Data availability

Original data generated and analyzed during this study are included in this published article or in supplementary materials. The RNAseq datasets were submitted to GEO with the series number GSE336666.

