## Supplementary material for "Type V Collagen Controls Decidual Extracellular Matrix Organization and Angiogenesis During Embryo Implantation": Gebril et al 2026 Supplementary File

### SUPPLEMENTARY METHODS

#### Superovulation

Three-week-old *Col5a1*$^{d/d}$ and *Col5a1*$^{f/f}$ mice were injected intraperitoneally with 5 IU pregnant mare serum gonadotropin (PMSG; Sigma-Aldrich). After 48 h, mice received 5 IU human chorionic gonadotropin (hCG; Sigma-Aldrich). At 16–18 h after hCG injection, mice were euthanized, and oocytes were collected from the oviductal ampulla and counted.

#### Measurement of serum estradiol and progesterone levels

Serum samples were collected on GD6 from *Col5a1*$^{f/f}$ and *Col5a1*$^{d/d}$ mice and submitted to the Ligand Assay and Analysis Core of the Center for Research in Reproduction at the University of Virginia for measurement of circulating 17β-estradiol (E2) and progesterone (P4) concentrations.

#### RNAscope

Paraffin-embedded tissues were sectioned, deparaffinized in xylene, and rehydrated in 100% ethanol. Endogenous peroxidase activity was blocked with RNAscope Hydrogen Peroxide (Cat. No. 322330, Advanced Cell Diagnostics) for 10 min at room temperature, followed by washing in distilled water. Target retrieval was performed using 1× target retrieval solution (Cat. No. 322000, Advanced Cell Diagnostics) for 15 min, followed by washes in distilled water and 100% ethanol. A hydrophobic barrier was drawn around each section, and slides were incubated with Protease Plus (Cat. No. 322330, Advanced Cell Diagnostics; 5 drops per slide) in a HybEZ oven (Boekel Scientific) at 40 °C for 30 min. Sections were then hybridized with a *Col5a1* probe (Cat. No. 521291, Advanced Cell Diagnostics; 4 drops per slide) at 40 °C for 2 h. Signal amplification was carried out using Amp 1–6 reagents (Cat. No. 322310, Advanced Cell Diagnostics) according to the manufacturer's protocol. Signals were detected using an equal mixture of BROWN-A and BROWN-B reagents (Cat. No. 322310, Advanced Cell Diagnostics). Sections were counterstained with hematoxylin for 2 min, washed in distilled water, dehydrated through 70% and 95% ethanol followed by xylene, and mounted with VectaMount (Vector Laboratories). Whole uterine sections were captured using the Leica VERSA8 whole slide imager and subsequently processed using Aperio ImageScope and ImageJ software.

### SUPPLEMENTARY FIGURES

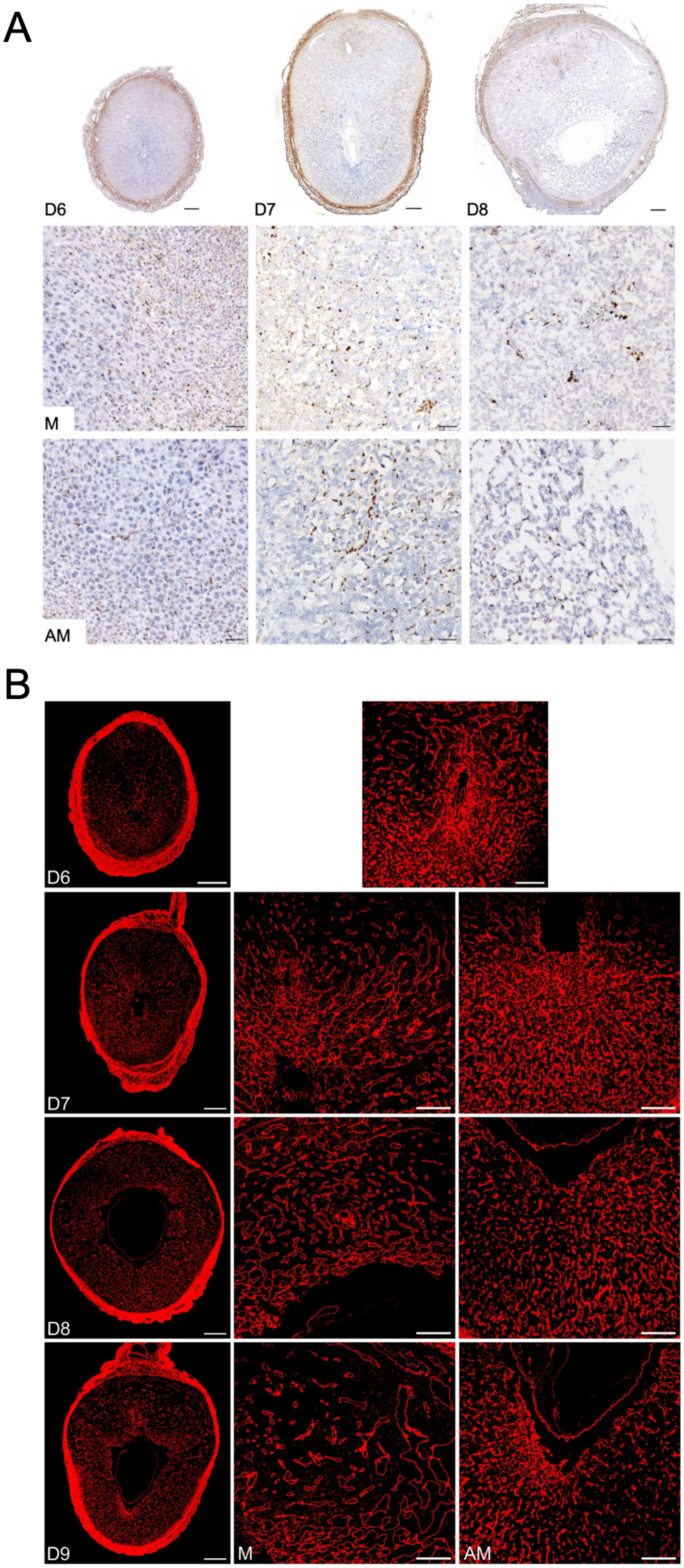

**Supplementary Fig. 1. Localization of *Col5a1* in the mouse uterus.**
(A) Localization of *Col5a1* mRNA in uterine sections from GD6 to GD8 mice using RNAscope (n = 3 per gestational time point). Scale bar: 250 µm (top panel) and 50 µm (middle and lower panels). Top panel: panoramic view of whole uterine sections; AM, images captured from the antimesometrial region; M, images captured from the mesometrial region.
(B) Confocal imaging of COL5A1 in frozen mouse uterine cross-sections from GD6 through GD9. Left panel: panoramic view of whole uterine sections; AM, images captured from the antimesometrial region; M, images captured from the mesometrial region. Imaging settings were adjusted for each individual image to optimize visualization of morphology. Representative images from three independent experiments. Scale bar: 500 µm (left panel) and 200 µm (all other images).

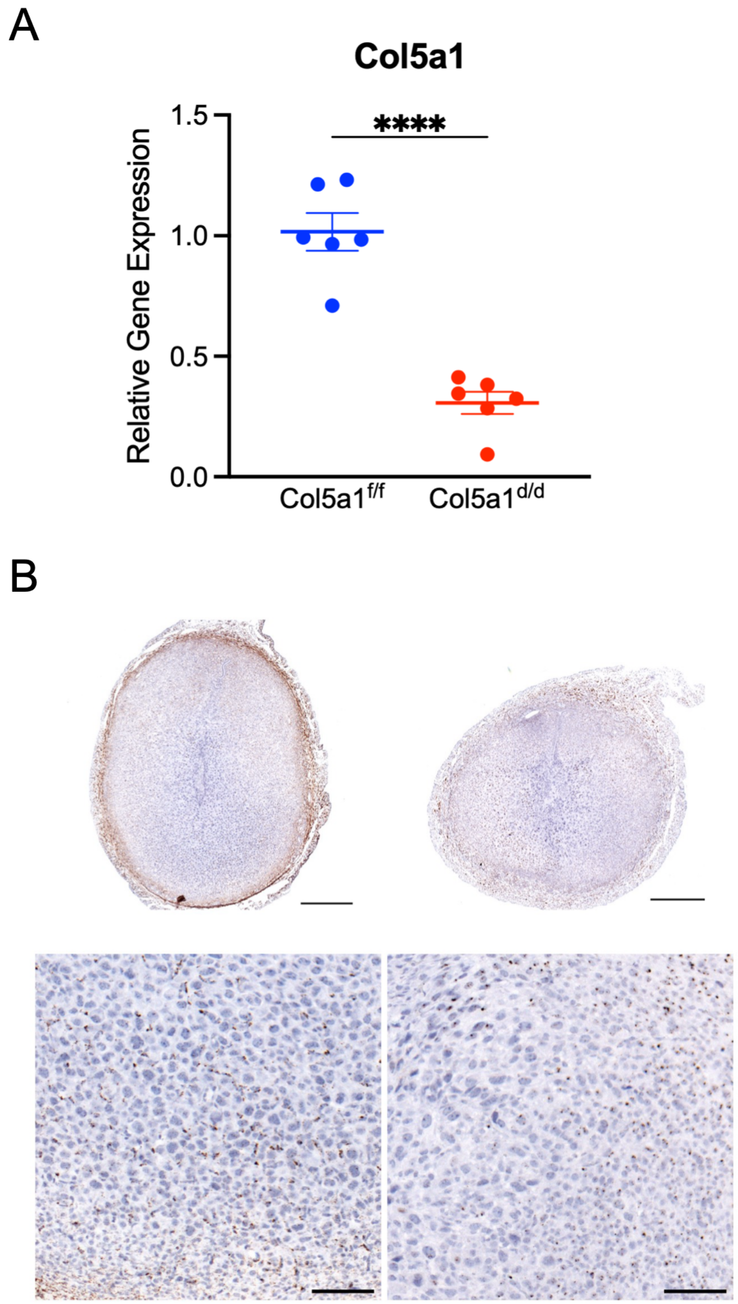


**Supplementary Fig. 2. Assessment of the efficiency of *Col5a1* deletion in the mouse decidua.**
(A) qPCR analysis of *Col5a1* gene expression in GD6 decidua from $Col5a1^{f/f}$ and $Col5a1^{d/d}$ mice (n = 6 per genotype; ****P < 0.0001).
(B) Localization of *Col5a1* mRNA in uterine sections from GD6 $Col5a1^{f/f}$ and $Col5a1^{d/d}$ mice (n = 3 per gestational time point). Scale bar: 500 µm (top panel) and 100 µm (lower panel).

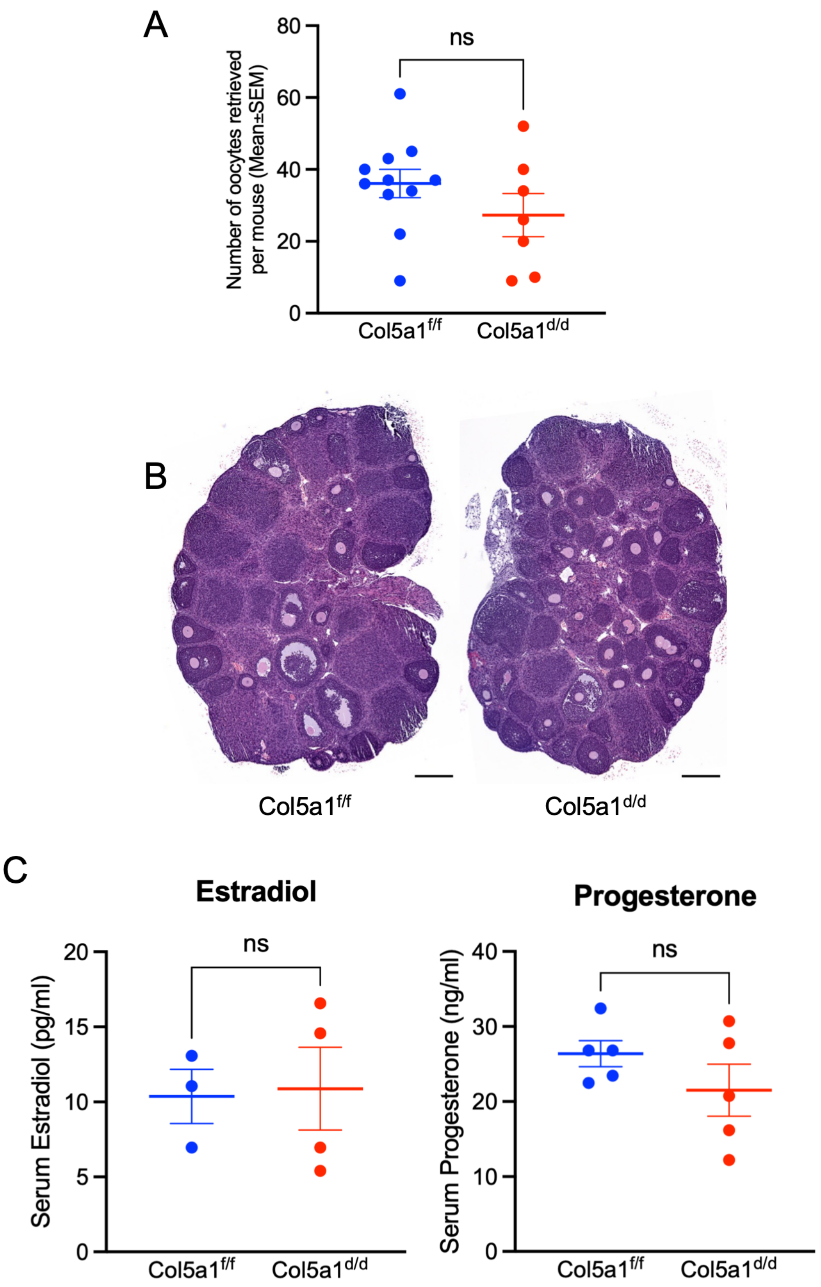


**Supplementary Fig. 3. Ovarian function is preserved in $Col5a1^{d/d}$ mice.**
(A) Dot plot showing the number of oocytes retrieved from the oviduct following superovulation, demonstrating comparable counts between groups ($Col5a1^{f/f}$: n = 11; $Col5a1^{d/d}$: n = 7). ns, not significant.
(B) Histological evaluation of ovaries from $Col5a1^{f/f}$ and $Col5a1^{d/d}$ mice following the superovulation procedure (n = 3 mice per genotype). Scale bar: 100 µm.
(C) Measurement of serum estradiol and progesterone levels at GD6 in $Col5a1^{f/f}$ and $Col5a1^{d/d}$ mice (n = 3–5 mice). ns, not significant.

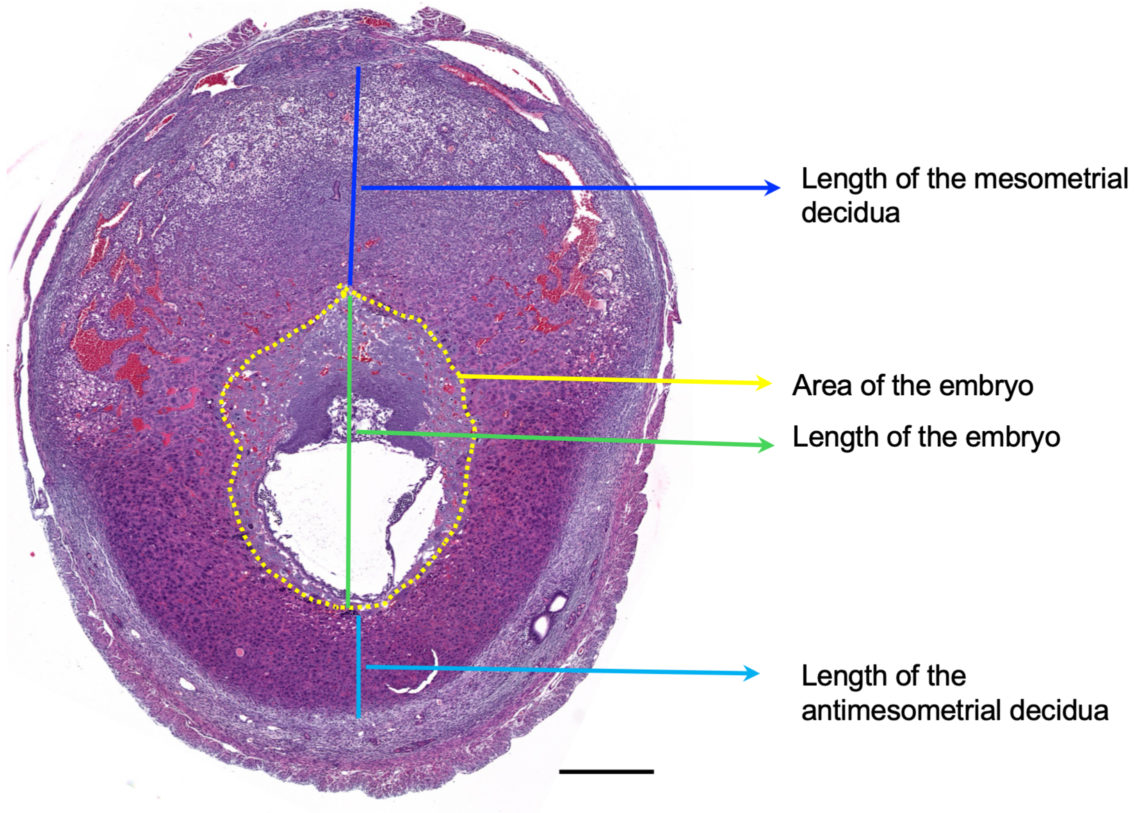


**Supplementary Fig. 4. Measurement guide for the decidua and embryo.**
Uterine sections were prepared from *Col5a1*$^{f/f}$ and *Col5a1*$^{d/d}$ mice at GD6–GD8 and stained with hematoxylin and eosin. Measurements, including the lengths of mesometrial and antimesometrial decidual regions, as well as embryo length and area, were obtained as outlined in the provided image using Aperio ImageScope software.

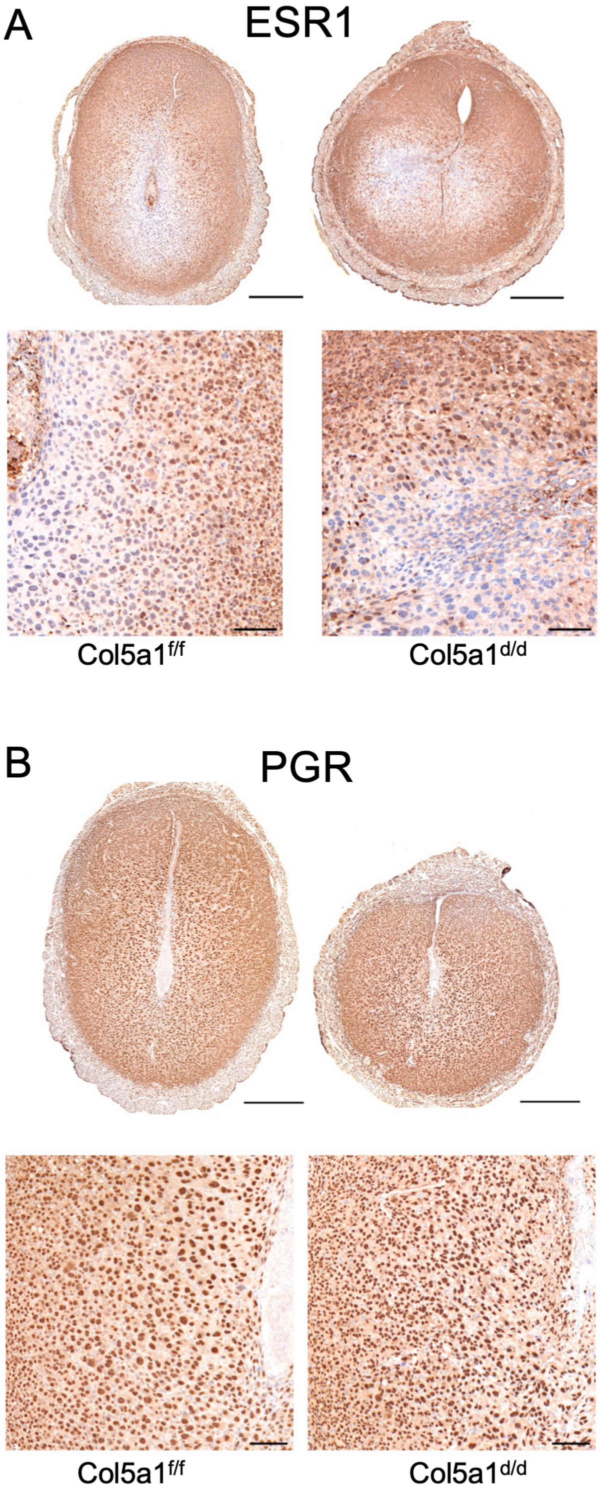


**Supplementary Fig. 5. Uterine ESR1 and PGR expression are not affected in *Col5a1$^{d/d}$* mice.**
(A) Immunohistochemical localization of ESR1 in GD6 uterine sections from *Col5a1$^{f/f}$* and *Col5a1$^{d/d}$* mice.
(B) Immunohistochemical localization of PGR in GD6 uterine sections from *Col5a1$^{f/f}$* and *Col5a1$^{d/d}$* mice.
Scale bar: 500 µm (top panel) and 100 µm (lower panel).

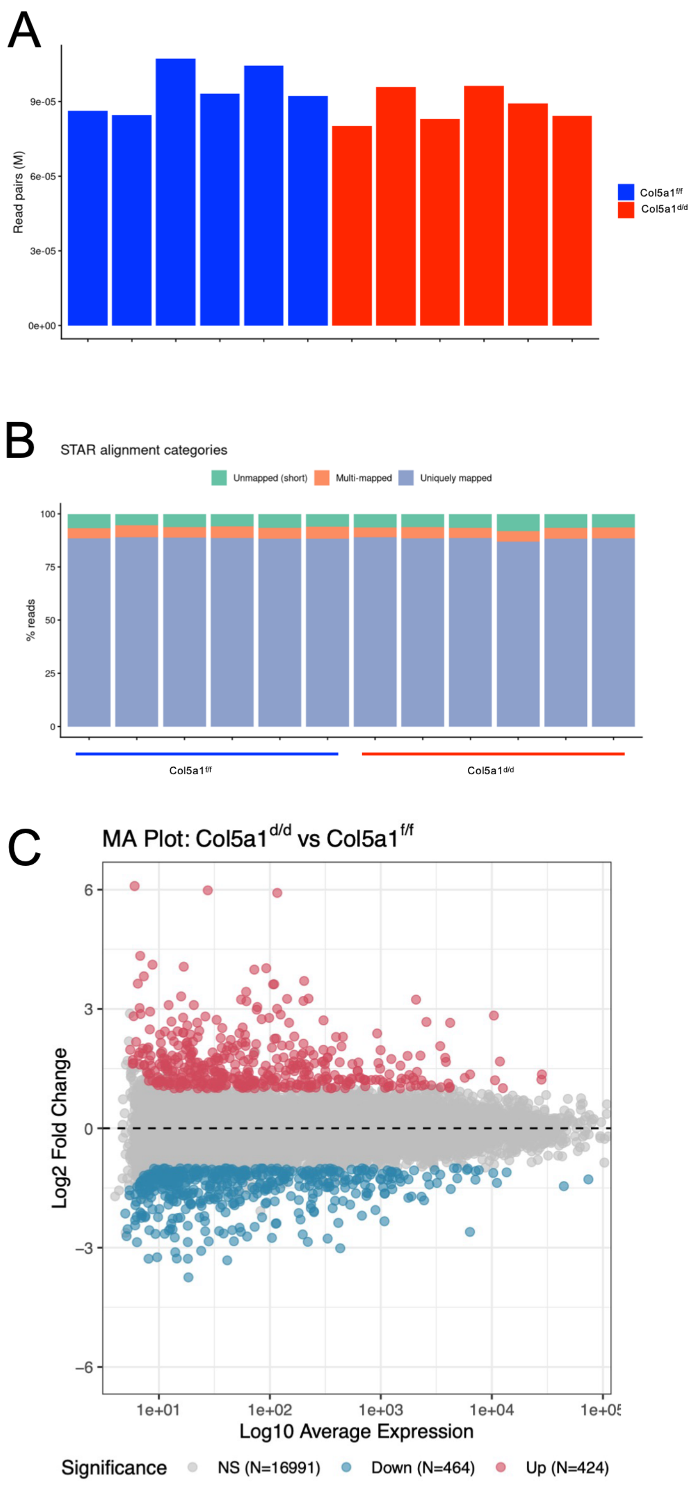


**Supplementary Fig. 6. Quality-control metrics from the nf-core/rnaseq pipeline.**
(A) Graph showing total read counts per sample generated by FastQC.
(B) Graph showing STAR alignment statistics for reads mapped to the reference genome, categorized as unmapped, multi-mapped, and uniquely mapped reads.
(C) MA plot showing log fold change relative to average abundance, where points distant from the horizontal center line indicate stronger condition-associated expression changes.

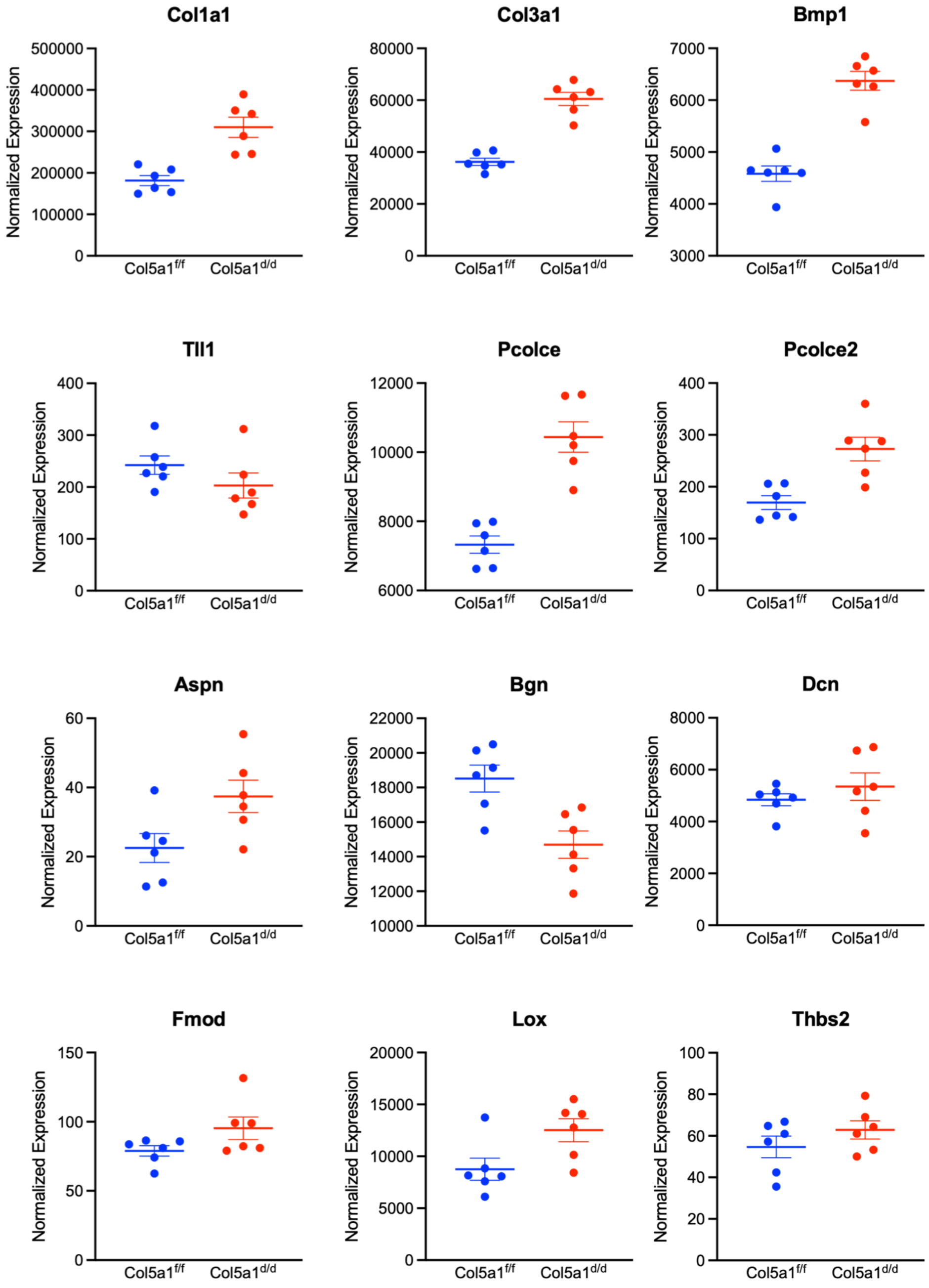


**Supplementary Fig. 7. Gene expression of factors involved in fibrillar collagen synthesis, processing, and assembly that are not altered in *Col5a1*$^{d/d}$ decidua.**

Normalized expression of genes in *Col5a1*$^{f/f}$ and *Col5a1*$^{d/d}$ decidua, as identified by RNA sequencing analysis. Expression of these genes is not significantly altered based on both thresholds (|$\log_2$ fold change| > 1 and adjusted P < 0.05).

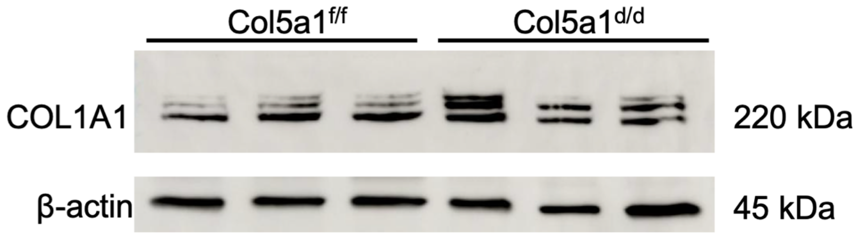


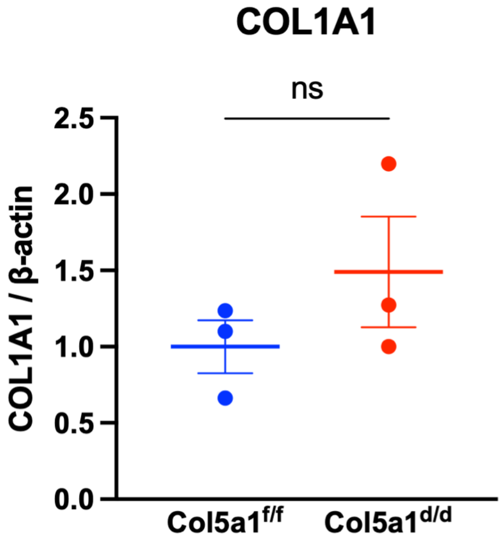


**Supplementary Fig. 8. COL1A1 production is not affected in *Col5a1*-deficient decidua.**
Western blot analysis of COL1A1 protein levels in GD6 decidua from *Col5a1*$^{f/f}$ and *Col5a1*$^{d/d}$ mice. Quantification of COL1A1 protein levels is shown in the graph.

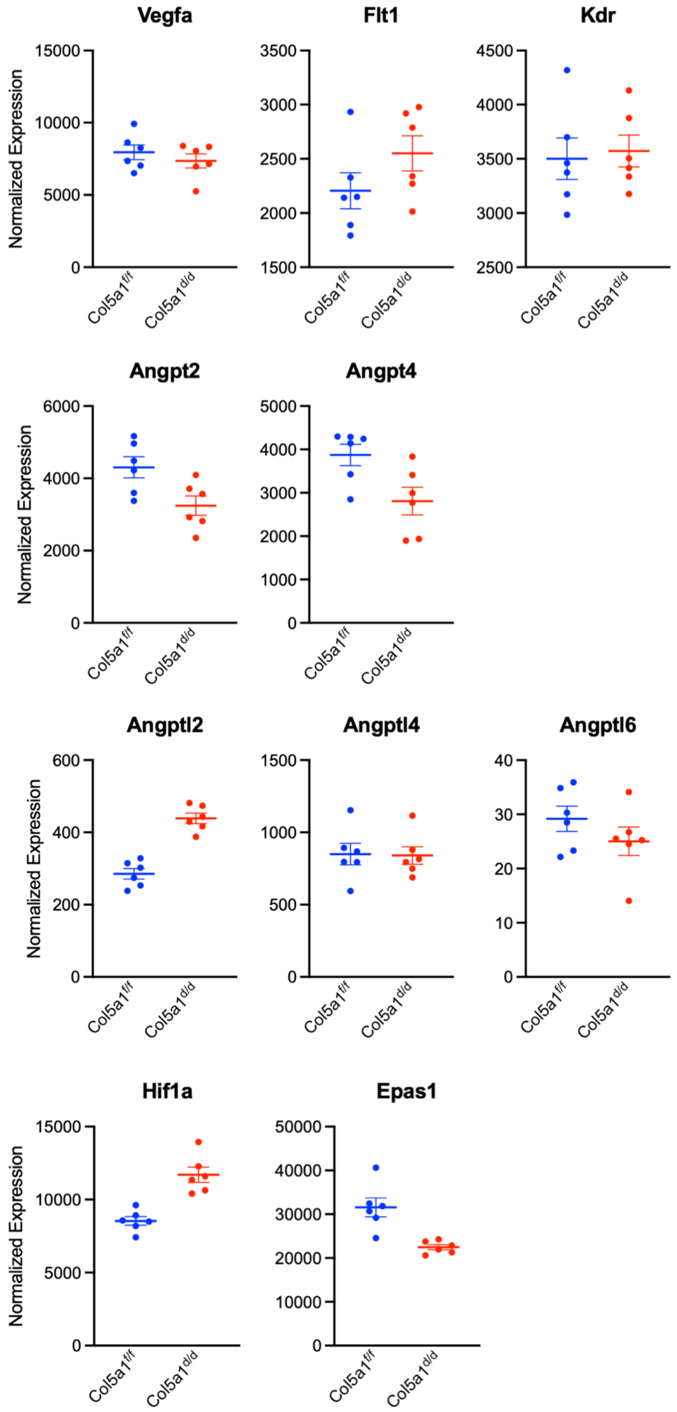


**Supplementary Fig. 9. Decidual angiogenesis markers that remain unaltered in *Col5a1*-deficient decidua.**

Normalized expression of genes in *Col5a1*$^{f/f}$ and *Col5a1*$^{d/d}$ decidua, as identified by RNA sequencing analysis. Expression of these genes is not significantly altered based on both thresholds ($|\log_2$ fold change$| > 1$ and adjusted $P < 0.05$).

**SUPPLEMENTARY TABLES**

**Supplementary table 1. Six months breeding experiment to evaluate reproductive phenotype.**

| Genotype | No. of mice | No. of litters weaned | No. of litters per mouse (Mean ± SEM) | No. of pups born | No. of pups per litter (Mean ± SEM) |
|---|---|---|---|---|---|
| $Col5a1^{f/f}$ | 7 | 37 | 5.3 ± 0.2 | 250 | 6.8 ± 0.4 |
| $Col5a1^{d/d}$ | 6 | 0 | 0 | 0 | 0 |

**Supplementary table 2. The total number of number of uteri examined per genotype with their total number of healthy and resorbed implantation sites counted at each gestational time point from Col5a1$^{f/f}$ and Col5a1$^{d/d}$ mice.**

| | **Col5a1$^{f/f}$** | | | **Col5a1$^{d/d}$** | | |
|---|---|---|---|---|---|---|
| | Total number of uteri examined | Total number of healthy implantation sites | Total number of resorbed implantation sites | Total number of uteri examined | Total number of healthy implantation sites | Total number of resorbed implantation sites |
| **Gestation Day 6** | 17 | 128 | 0 | 18 | 129 | 0 |
| **Gestation Day 7** | 15 | 115 | 0 | 21 | 145 | 0 |
| **Gestation Day 8** | 17 | 129 | 0 | 25 | 136 | 28 |
| **Gestation Day 9** | 6 | 41 | 0 | 10 | 10 | 54 |
| **Gestation Day 10** | 4 | 33 | 0 | 4 | 0 | 30 |
| **Gestation Day 12** | 3 | 20 | 0 | 3 | 0 | 20 |

**Supplementary table 3. The list of primers used in this study**

| Oligo name | Forward | Reverse |
|---|---|---|
| **Col5a1** | GCTGAAAATACACACAAAGTAAACAGAA | TGTACCGTTTGGCCTGGAAT |
| **Prl8a2** | AAACCCACCAGCTCATGGAC | GGAGTGATCCATGCACCCAT |
| **Bmp2** | CAAAGCAGGACCAGTGGGAA | AGCCCCCTGGAAGGGATTAT |
| **Wnt4** | GTACCTGGCCAAGCTGTCAT | CCGGAACTGGTATTGGCACT |
| **Gja1** | TGGACAAGGTCCAAGCCTACTC | TCCCCAGGAGCAGGATTC T |
| **Rplpo** | CACTGGTCTAGGACCCGAGAAG | GGTGCCTCTGGAGATTTTCG |
